# Competing Subclones and Fitness Diversity Shape Tumor Evolution Across Cancer Types

**DOI:** 10.1101/2025.05.31.657191

**Authors:** Hai Chen, Jingmin Shu, Rekha Mudappathi, Elaine Li, Panwen Wang, Leif Bergsagel, Ping Yang, Zhifu Sun, Logan Zhao, Changxin Shi, Jeffrey P. Townsend, Carlo Maley, Li Liu

## Abstract

Intratumor heterogeneity arises from ongoing somatic evolution complicating cancer diagnosis, prognosis, and treatment. Here we present TEATIME (es*t*imating *<u>e</u>*volution*<u>a</u>*ry events *t*hrough s*i*ngle-ti*<u>m</u>*epoint s*<u>e</u>*quencing), a novel computational framework that models tumors as mixtures of two competing cell populations: an ancestral clone with baseline fitness and a derived subclone with elevated fitness. Using cross-sectional bulk sequencing data, TEATIME estimates mutation rates, timing of subclone emergence, relative fitness, and number of generations of growth. To quantify intratumor fitness asymmetries, we introduce a novel metric—fitness diversity—which captures the imbalance between competing cell populations and serves as a measure of functional intratumor heterogeneity. Applying TEATIME to 33 tumor types from The Cancer Genome Atlas, we revealed divergent as well as convergent evolutionary patterns. Notably, we found that immune-hot microenvironments constraint subclonal expansion and limit fitness diversity. Moreover, we detected temporal dependencies in mutation acquisition, where early driver mutations in ancestral clones epistatically shape the fitness landscape, predisposing specific subclones to selective advantages. These findings underscore the importance of intratumor competition and tumor-microenvironment interactions in shaping evolutionary trajectories, driving intratumor heterogeneity. Lastly, we demonstrate that TEATIME-derived evolutionary parameters and fitness diversity offer novel prognostic insights across multiple cancer types.

## INTRODUCTION

A tumor is composed of diverse cell populations, each with distinct molecular and phenotypic profiles^1,2^. This diversity enables a tumor to adapt to various selection pressures, such as changes in the microenvironment and treatment, leading to proliferation of subpopulations with greater fitness^3,4^. Understanding the evolutionary parameters governing proliferation and survival of these subpopulations is crucial for gaining insights into tumor development, metastasis, therapy resistance, and disease recurrence.

The clonal evolution model posits that a tumor originates from a single cell that has acquired genetic or epigenetic alterations conferring a growth advantage. As this cell proliferates, it forms a population (i.e., clone) that can further evolve by accumulating new alterations. While most of these alterations are selectively neutral, some provide additional growth advantages^5^, leading to the emergence of subpopulations (i.e., subclones) with increased fitness^6,7^. This evolutionary process continuously shapes the cellular composition of a tumor, resulting in complex intratumor heterogeneity.

Tumor evolutionary processes are often reconstructed by analyzing somatically acquired single nucleotide alterations (SNAs), owing to their high abundance, close adherence to the infinite-site mutation model, and relatively stable mutation rates. Although SNAs do not account for all tumorigenic events, they constitute many key driver alterations and may serve as a reliable proxy. Qualitative analyses of tumor evolution typically categorize SNAs into early vs. late, or clonal vs. subclonal groups^8–10^. This temporal order has been associated with molecular and phenotypic traits of tumors. Quantitative approaches estimate key evolutionary parameters such as mutation rates and relative fitness of subclones, and cellular composition metrics, offering a more dynamic and mechanistic understanding of tumor development. Williams et al. pioneered a mathematical model of clonal evolution to estimate evolutionary parameters using variant allele frequencies (VAFs) obtained from whole-exome sequencing (WES) or whole-genome sequencing (WGS) data^11^. Building on this foundation, several subsequent studies have explored machine learning and other techniques to enhance the estimations^12–14^. However, these analyses have relied on one or both of the following assumptions: first, the tumor cell population size at the time of sampling is known; and second, the number and VAF of low-frequency variants follow a power-law distribution^11^, although these variants are polyphyletic and do not share a common trajectory. These simplifications can lead to model convergence failures in many tumors, compromise the accuracy of the estimates, and limit the biological relevance of the results.

To address these limitations, we developed a method named es*t*imating *<u>e</u>*volution*<u>a</u>*ry events *t*hrough s*i*ngle-ti*<u>m</u>*epoint s*<u>e</u>*quencing (TEATIME). This method models the tumor cell population size as a dynamic variable, tracks the temporal changes of VAFs as the tumor evolves, and jointly infers mutation clusters and evolutionary parameters in an iterative framework. Extensive simulations demonstrated that TEATIME enhanced the accuracy and robustness of parameter estimation generally. When applied to WES data from 33 diverse tumor types cataloged in The Cancer Genome Atlas (TCGA), TEATIME increased the number of analyzable tumors by 2– to 37–fold compared to existing methods. TEATIME also revealed striking divergences and conserved patterns in evolutionary profiles. Furthermore, we introduce a novel measure, fitness diversity, that captures the imbalance between subclones with distinct fitness levels. Fitness diversity represents a new conceptual dimension regarding intratumor heterogeneity. By linking quantitative evolutionary parameters with immune context, pathway activity, and patient prognosis, TEATIME provides a unified framework enabling understanding of key properties of tumor evolution, as well as offering a foundation for evolution-informed therapeutic strategies.

## RESULTS

### TEATIME algorithm

We model cell division and mutation accumulation in a tumor as a discrete-time, discrete-state Markov process ^13–15^ (**Fig. 1A**). Starting with a single cell at time point *t*_0_, its mitosis produces two daughter cells. Each daughter cell has a probability of dying before entering the next division cycle. Continued division of surviving cells over time leads to the formation of a population, with the initial cell at *t*_0_ being the most recent common ancestor (*MRCA*_0_). The growth of this population follows an exponential model, such that at a given time point *t* > *t*_0_, the number of cells descended from the *MRCA*_0_ is

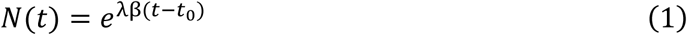

where λβ is the cell proliferation rate (**Methods and Materials**). The unit of time is defined as the time between two consecutive cell divisions, so *t* can be interpreted as the number of generations. In a successful cell division, somatic mutations present in the parent cell are passed to the daughter cells and additional mutations are introduced at a rate μ, which is the number of mutations acquired per diploid genome per cell division.

**Figure 1.**
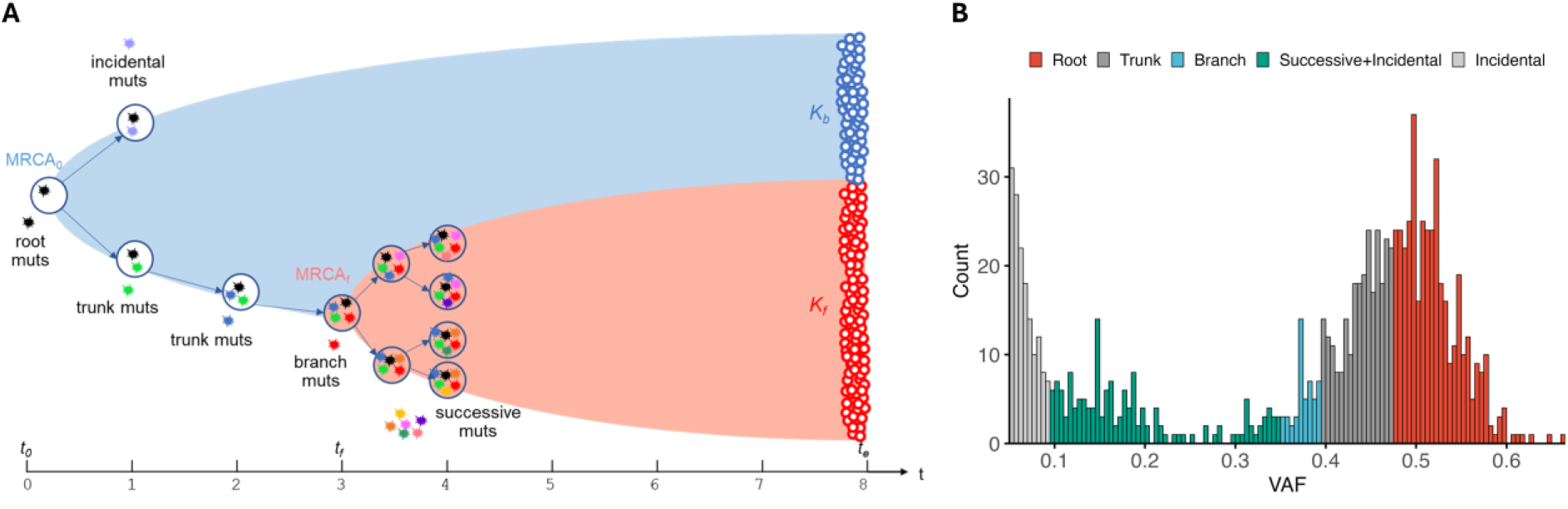
Schematic representation of TEATIME algorithm. **(A)** Tumor growth modeled as a discrete-time discrete-state Markov process. Cell divisions occur at a constant rate and at specific time points. Each division introduces mutations. The state of a cell is defined by its mutation composition. Asterisks with different colors represent mutations acquired at different time points. At time *t*_0_, a single cell initiates the growth of a cell population via division. The growth progresses to the emergence of a subclone at time *t*_*f*_, and continues until the point of sampling and sequencing at time *t*_*e*_. Among cells with fitness similar to that of the initial cell (blue circles), 1 division cycle occurs in one unit of time.

If all genetic or epigenetic alterations occurred after *t*_0_ are selectively neutral, the cells grow at the same rate of λβ and form the ancestral clone K_*a*_. If a cell acquires alterations at time *t*_*f*_, which confer a growth advantage, it becomes the MRCA_*f*_ and proliferates at a rate λβ(1 + s) with s > 0, forming a subclone K_*f*_. The fitness advantage of K_*f*_ over K_*a*_ is quantified by the selection coefficient s, defined as the relative increase in net growth rates between the two populations. At the time point *t*_*e*_ when a tumor is sampled for analysis, the total number of cells is *N*. The fraction of cells belonging to subclone K_*f*_ among all tumor cells at *t*_*e*_ is

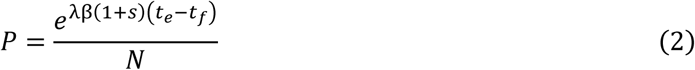

The fraction of cells in K_*f*_ over those in K_*a*_ is determined by the evolutionary parameters as

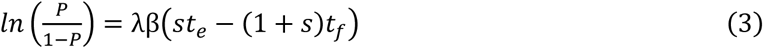

with a higher fitness than the initial cell (pink circles), >1 division cycle occurs in one unit of time. Blue and pink areas represent the expansions of the background clone K_*b*_ and the advantageous subclone K_*f*_, respectively. **(B)** Decomposing the VAF distribution to different mutation groups.

Among cells Mutations acquired in the “foundation lineage” represent the evolutionary path from *MRCA*_0_ to MRCA_*f*_ (Fig. 1A). This lineage includes three classes of mutations depending on the time of acquisition. Root mutations, which preexist at *t*_0_, are present in all tumor cells. Trunk mutations, acquired after *t*_0_ but before *t*_*f*_, are present in all K_*f*_ cells and some K_*a*_ cells. For a trunk mutation acquired at time *t*, the fraction of tumor cells carrying it at *t*_*e*_ is *f*_*k*_(*t*). The number of trunk mutations accumulated till time *t* is

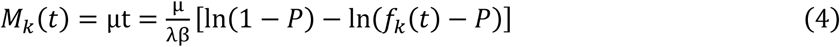

A total of μ branch mutations are acquired by the MRCA_*f*_ cell at time *t*_*f*_. They are present in all K_*f*_ cells but absent in K_*a*_ cells. The fraction of cells carrying branch mutations at *t*_*e*_ is *f*_ℎ_ = *P*.

Outside the foundation lineage there are incidental mutations that are present in some K_*a*_ cells but absent in K_*f*_ cells. There are also successive mutations that are present in some K_*f*_ cells but absent in K_*a*_ cells. For a successive mutation acquired at time *t*, the fraction of cells carrying it at *t*_*e*_ is

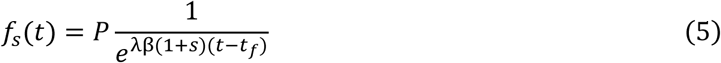

The above equations link the evolutionary process of a tumor to the fractions of cells carrying specific mutations at the time of sampling. These fractions can be inferred from VAFs reported by WES or WGS of bulk tumor cells^58^. For a heterozygous mutation in a diploid region, its VAF corresponds to half the fraction of cells carrying this mutation after adjusting for sample impurity. Leveraging these relationships, TEATIME transforms the estimation of evolutionary parameters to a task of decomposing a tumor’s mutational compositions (Fig. 1B). By maximizing the likelihood of observing the number of mutations in various categories and their VAF distributions, TEATIME infers maximum likelihood estimates for a set of parameters, including μ, *s, t*_*f*_, and *P* (**Supplementary Materials**).

### Evaluation of TEATIME performance

We used simulation data to evaluate the performance of TEATIME, MOBSTER, and TumE, all of which operate in the subclonal evolution framework. The first simulated dataset, included with the MOBSTER package, comprises 150 synthetic tumor samples generated to follow a subclonal evolution trajectory based on the discrete-time discrete-state Markov process ^16^. In these simulations, the following parameters were held constant, including *N* = 10^8^, μ = 16, sequencing depth at 120× and a cell death rate of 0.2. Other parameters were varied across defined ranges, including *t*_*f*_ ∈ [4, 14], *s* ∈ [0.125, 1.625], and *P* ∈ [0, 0.97].

These tools did not produce results for all simulated tumors. We considered a tumor non-analyzable if the TEATIME or MOBSTER model failed to infer the complete set of evolutionary parameters. For TumE, tumors with fewer than five valid Monte Carlo estimates were deemed non-analyzable. MOBSTER was able to analyze only 42% of the simulated tumors, whereas TumE and TEATIME achieved success rates exceeding 60% and 70%, respectively (Fig. 2A). Notably, the non-analyzable tumors tended to have more extreme subclone fractions, either very low or very high, compared to the subclone fractions observed in analyzable tumors (Fig. 2B). To ensure a fair comparison, we restricted the subsequent analyses to the subset of 37 tumors that were successfully analyzed by all three methods. Since TumE produced a wide range of values for each parameter due to its Monte Carlo sampling approach (**Supplementary** Fig. 3A-B), we used the mean as the final estimation.

**Figure 2.**
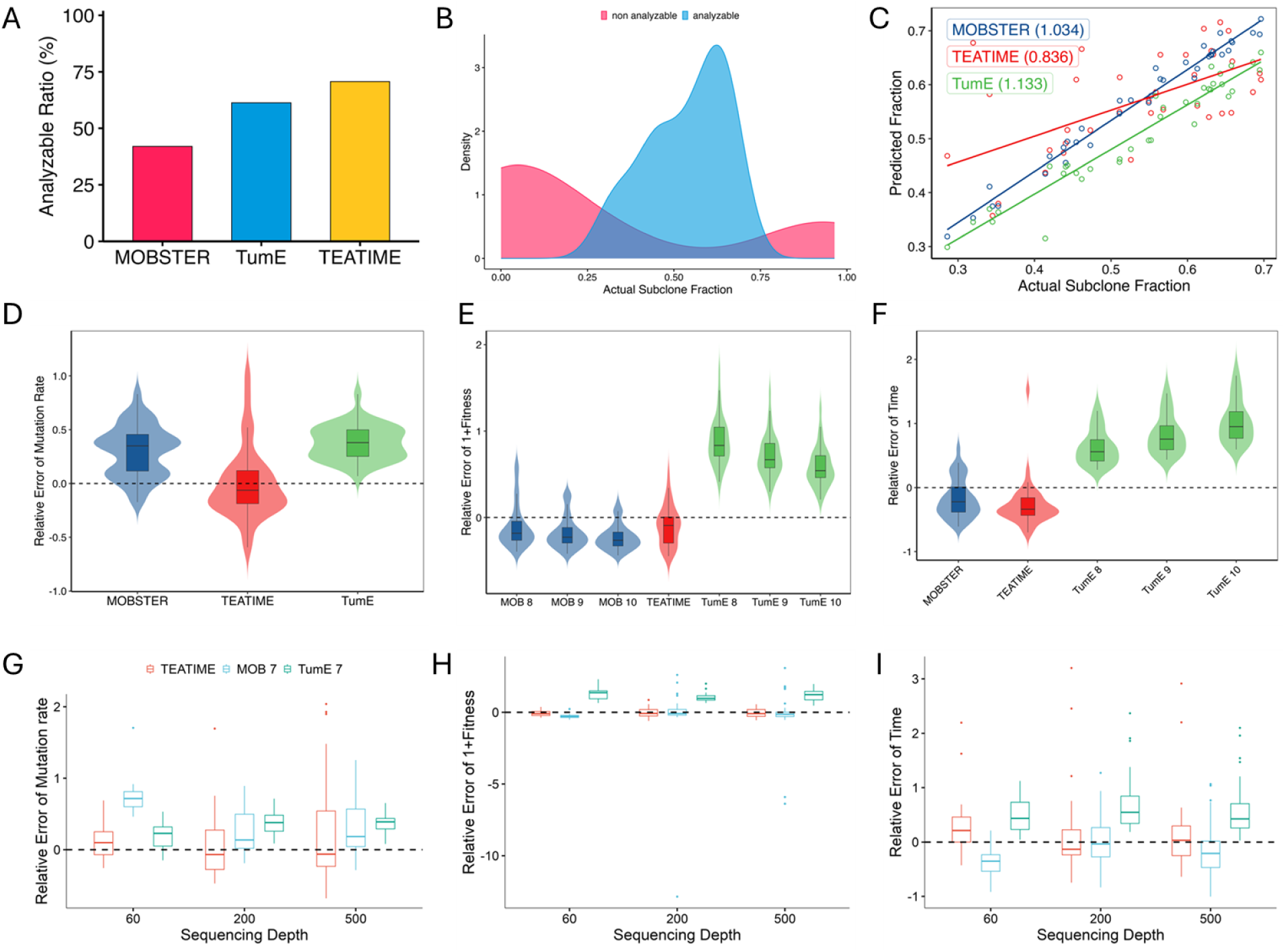
Comparison of the TEATIME, MOBSTER, and TumE on simulation datasets. (**A**) Proportion of analyzable tumors. (**B**) Density plot shows distributions of subclone fraction in tumors analyzable versus non-analyzable. (**C**) Scatter plot shows correlation between estimated and simulated subclone fractions. All regression slopes are highly significant (*P* < 0.001). (**D-F**) Violin plots show distributions of relative errors of estimated mutation rate (μ), fitness (s), and time of subclone emergence (*t*_*f*_). A range of *N* values from 10^8^ to 10^10^ were provided to MOBSTER and TumE as input. Each configuration is annotated accordingly (e.g., MOB 8 denotes MOBSTER with *N* =10^8^). (**G-I**) comparison on datasets with varying sequencing depth and *N =*10^7^.

Despite having the lowest proportion of analyzable tumors, MOBSTER’s estimated subclone fractions showed an almost perfect correlation with the simulated values (Regression coefficient RC = 1.03, Fig. 2C). TEATIME and TumE also demonstrated high accuracy, with slightly lower but still very strong correlations (RC = 0.84 and 1.13, respectively). When evaluating additional parameters, including *μ, s* and *t*_*f*_, TEATIME consistently outperformed the other two methods, achieving the lowest relative errors centered around zero (Fig. 2D**-F**). In contrast, MOBSTER tended to overestimate *μ* and underestimate *s* and *t*_*f*_, whereas TumE systematically overestimated all three parameters (Fig. 2D**-F**). Because MOBSTER and TumE require *N* to calculate *s* or *t*_*f*_, we examined the sensitivity of their estimations to different values of *N* (**Supplementary** Fig. 3C-D). Surprisingly, providing a value of *N* that deviated from the simulated ground truth sometime led to smaller errors.

To complement the initial simulations where sequencing depth and final cell population size were fixed, we used the TEMULATOR ^16^ to produce additional datasets using a series of *d* ranging from 60× to 1000× and *N* ranging from 10^6^ to 10^7^. Interestingly, increasing sequencing depth and providing true N values did not improve the estimations for MOBSTER and TumE (Fig. 2G**-I****, Supplementary** Fig. 4-5). In contrast, TEATIME produced parameter estimates with relative errors more tightly centered around zero, particularly at sequencing depths exceeding 100×. Because TumE outputs multiple values for each parameter, we plotted the 95% confidence ellipses of relative errors using all Monte Carlo estimates alongside TEATIME predictions. The ellipse area for TumE was markedly larger than that of TEATIME. Again, increasing sequencing depth did not substantially reduce the ellipse area for TumE, in contrast to TEATIME, where higher depth led to more accurate estimates (**Supplementary** Fig. 4D**, 5D**).

These results collectively demonstrated that TEATIME outperformed MOBSTER and TumE by producing minimal expected relative error in estimating *μ, s*, and *t*_*f*_ without requiring prior information on *N*.

### Pan-cancer Analysis of Tumor Evolution

We extracted WES data of 8,935 primary tumors representing 33 cancer types from the TCGA study ^17^. For each tumor, we used SNVs located in diploid genomic regions to infer evolutionary parameters. Because less than 8% of these tumors were analyzable by MOBSTER and TumE (**Supplementary** Fig. 6), we focused on TEATIME analysis results.

TEATIME produced evolution estimators for 1,628 (18%) tumors (Fig. 3A**, Supplementary Table 1**). The proportion of analyzable tumors varied substantially across cancer types. For example, in colon adenocarcinoma (COAD), approximately 40% of tumors yielded results, whereas in acute myeloid leukemia (AML), only a single case was analyzable. Because TEATIME assumes the presence of two subpopulations with distinct fitness levels, tumors deemed non-analyzable likely exhibit a more homogeneous fitness landscape compared to those that are analyzable. This hypothesis aligns with the stem cell model of AML, in which a small population of cancer stem cells continuously gives rise to progenitor cells that differentiate into tumor cells without forming an apparent subclonal structure ^18,19^. To ensure sufficient statistical power, we limited the subsequent analysis to 19 tumor types that had at least 20 analyzable tumors.

**Figure 3.**
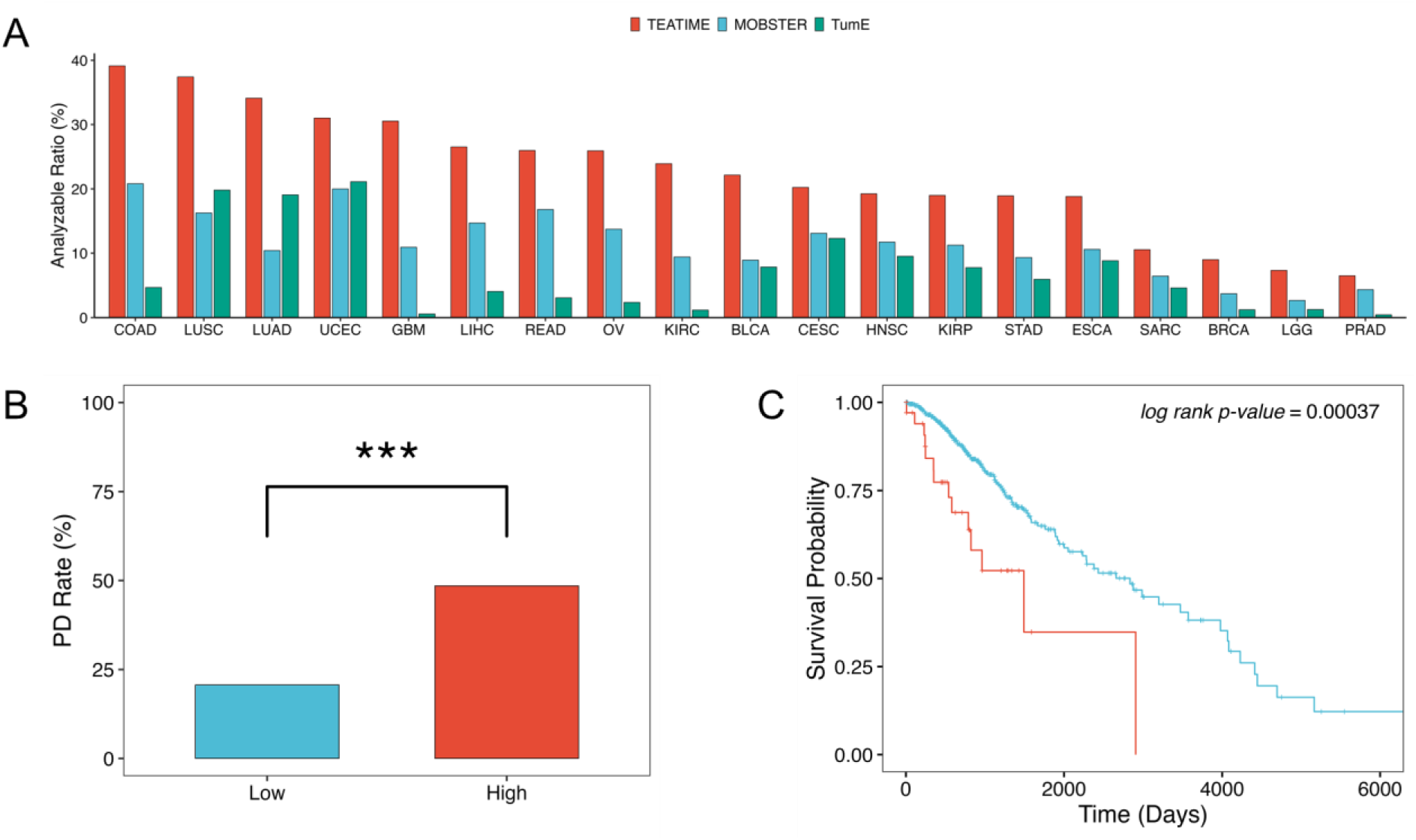
Fitness diversity across cancer types and its clinical relevance. (**A**) Fraction of analyzable tumors by cancer type. (**B**) Fraction of tumors exhibiting progressive disease (PD) after treatment, stratified by median fitness index θ (red: high θ; blue: low θ). *** indicates p < 0.001. (**C**) Kaplan–Meier plot comparing overall survival between low-θ and high-θ groups (log-rank p < 0.001).

### Fitness diversity is a new dimension of intra-tumor heterogeneity

The extent of mixing between K_*a*_ and K_*f*_ clones in a tumor reflects the degree of heterogeneity among competing cell populations. To quantify this, we define a fitness diversity index θ based on Shannon entropy, which captures the imbalance between cell populations with distinct fitness levels (**Methods and Materials**). The θ ranges from 0 to 1, with 0 indicating a tumor composed entirely of K_*a*_ or K_*f*_ clones (i.e., no diversity), and 1 indicating a perfectly balanced mixture of K_*a*_ and K_*f*_ clones. For non-analyzable tumors, we assume they are dominated by either K_*a*_ or K_*f*_, and assign θ = 0. We found that θ is a novel prognostic marker. In low-grade glioma (LGG), tumors with high θ were associated with an increased risk of poor response to treatment (*P* =8×10^-5^, Fig. 3B) and shorter overall survival (*P* =4×10^-4^, Fig. 3C). This association remained significant even after adjusting for age, sex and *IDH1* mutation status (Cox regression *P* =9×10^-4^), suggesting θ is an independent prognostic marker.

### Pan-cancer subclonal evolution characteristics

Mutation rates varied across cancer types. On average, the mutation rate of LGG tumors was the lowest (mean *μ* =2.9 per exome per generation), while uterine corpus endometrial carcinoma (UCEC) was highest (mean *μ* =8.9 per exome per generation, Fig. 4A)^20^. In most tumor types, derived subclones emerged early. The mean *t*_*f*_ ranged from 1 generation after *t*_0_ in LGG to 7 generations in UCEC (Fig. 4B). This observation is consistent with previous studies reporting that aggressive subclones, such as those associated with metastasis or treatment resistance, often arise near the onset of tumorigenesis ^21–23^. To quantify the duration of subclone expansion relative to its emergence, we calculated a subclone expansion score τ = *t*_*e*_/*t*_*f*_. This score ranged from a mean of 4.4 in esophageal carcinoma (ESCA) to 7.9 in bladder urothelial carcinoma (BLCA, Fig. 4C), suggesting that most subclones undergo extended growth prior to clinical detection. Meanwhile, the estimated selection coefficients *s* are moderate across all tumors, with none exceeding 2.04 (Fig. 4D). These findings indicate that despite the sustained expansion of subclones, their modest fitness advantages constrain rapid or explosive growth.

**Figure 4.**
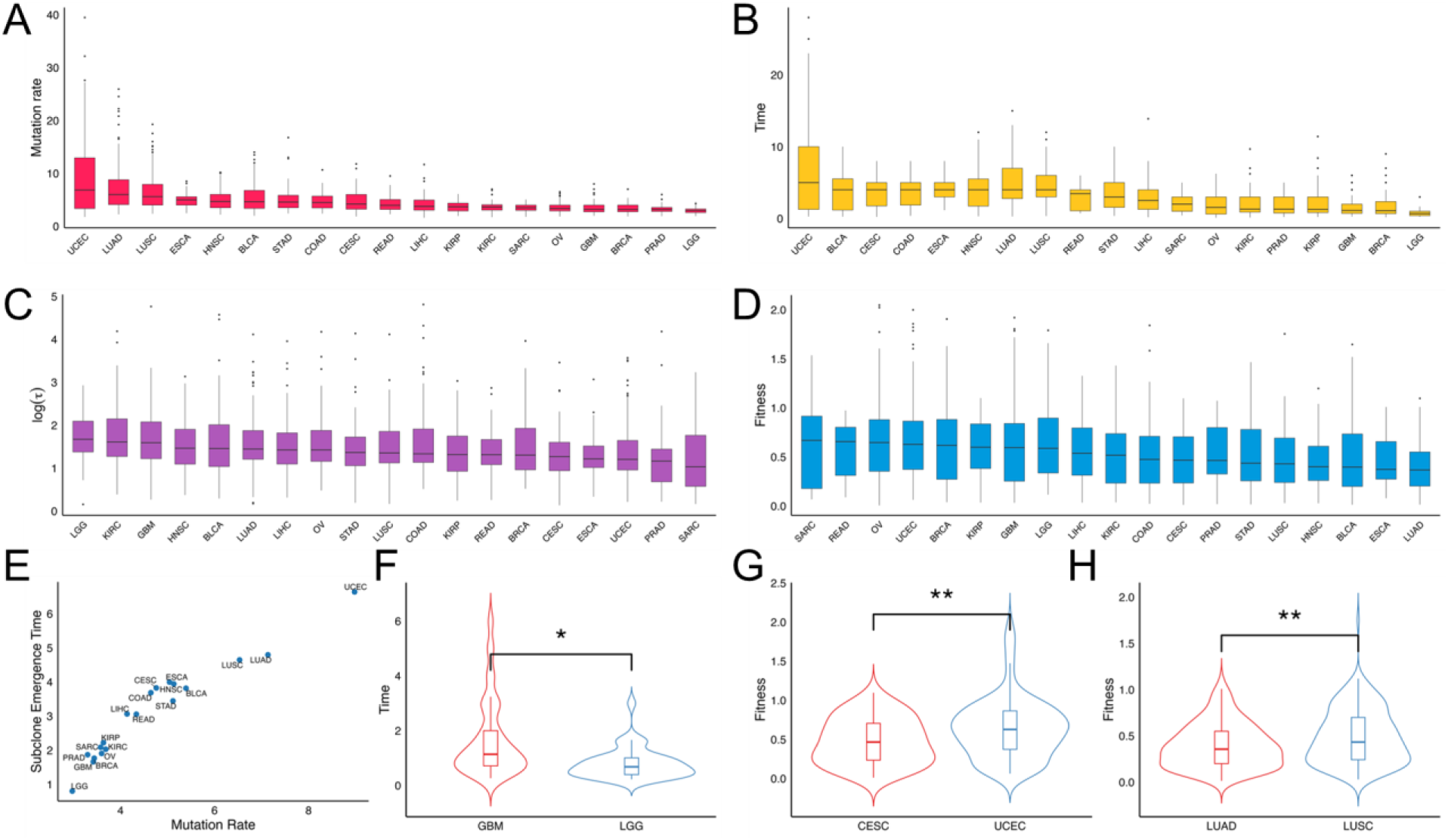
Pan-cancer distribution of inferred evolutionary parameters. (**A-D**) Boxplots show distributions of mutation rate (A), subclone emergence time (B), subclone expansion score (C) and subclone fitness (D) across cancer types. (**E**) Cancer types with higher mean mutation rates tend to exhibit later subclone emergence time. (**F-H**) Violin plots show distributions of inferred evolutionary parameters between cancer types in the same organ: brain (F), uterus (G), and lung (H).

Interestingly, we found that in tumor types with high mutation rates, derived subclones tended to emerge relatively late (Fig. 4E). These tumors, including UCEC, lung adenocarcinoma (LUAD), lung squamous cell carcinoma (LUSC), BLCA, and ESCA, are known to exhibit high mutational burdens, often resulting from prolonged exposure to inflammation, environmental insults, or other factors during extended precancerous stages^24–26^. The delayed emergence of subclones in these contexts suggests that while mutational accumulation occurs early and broadly, the selective advantage required for subclonal expansion may only arise later, possibly due to microenvironmental shifts or late-arising driver mutations.

Cancers arising in the same organ but differing in histological type often display distinct clinical features. For example, LGG typically presents at a younger age than GBM ^27^, and LUSC is associated with larger tumor size and poorer outcomes compared to LUAD ^28–30^. Consistent with these clinical distinctions, we observed significant differences in their evolutionary dynamics. In brain tumors, LGG exhibited an earlier subclone emergence time than glioblastoma multiforme (GBM, *P* =0.02, Fig. 4F). In uterine tumors, subclone fitness was higher in UCEC compared to cervical squamous cell carcinoma and endocervical adenocarcinoma (CESC, *P* =0.0044, Fig. 4G). A similar trend was observed in lung cancers, where subclones in LUSC exhibited markedly higher fitness than those in LUAD (*P* =0.002, Fig. 4H). These findings warrant further investigation into how evolutionary differences relate to underlying clinical features.

### Association of immune infiltration with tumor heterogeneity and evolutionary trajectory

In addition to intratumor competition, the immune microenvironment may also impose selective pressures to shape a tumor’s evolutionary trajectory. To investigate this, we obtained pre-computed tumor-infiltrating immune cell abundances in TCGA tumors using three tools, including TIMER, xCell, and CIBERSORT^31–33^. We then performed a pan-cancer multivariate regression analysis to assess associations between immune cell infiltration scores and evolutionary parameters (**Methods & Materials**). Our analysis revealed that both the fitness diversity index θ and subclone selection coefficient *s* were consistently and negatively associated with the infiltration levels of multiple immune cell types, while subclone emergence time *t*_*f*_ showed positive associations (FDR<0.05, **Fig 5A**). These patterns suggest that in tumors with abundant immune infiltration, subclones tend to emerge later and exhibit lower selection advantages, resulting in reduced fitness diversity. While previous studies have shown that “immune-hot” microenvironments can constrain intratumor genetic heterogeneity heterogeneity^34–40^, our results offer additional insight from an evolutionary perspective.

**Figure 5.**
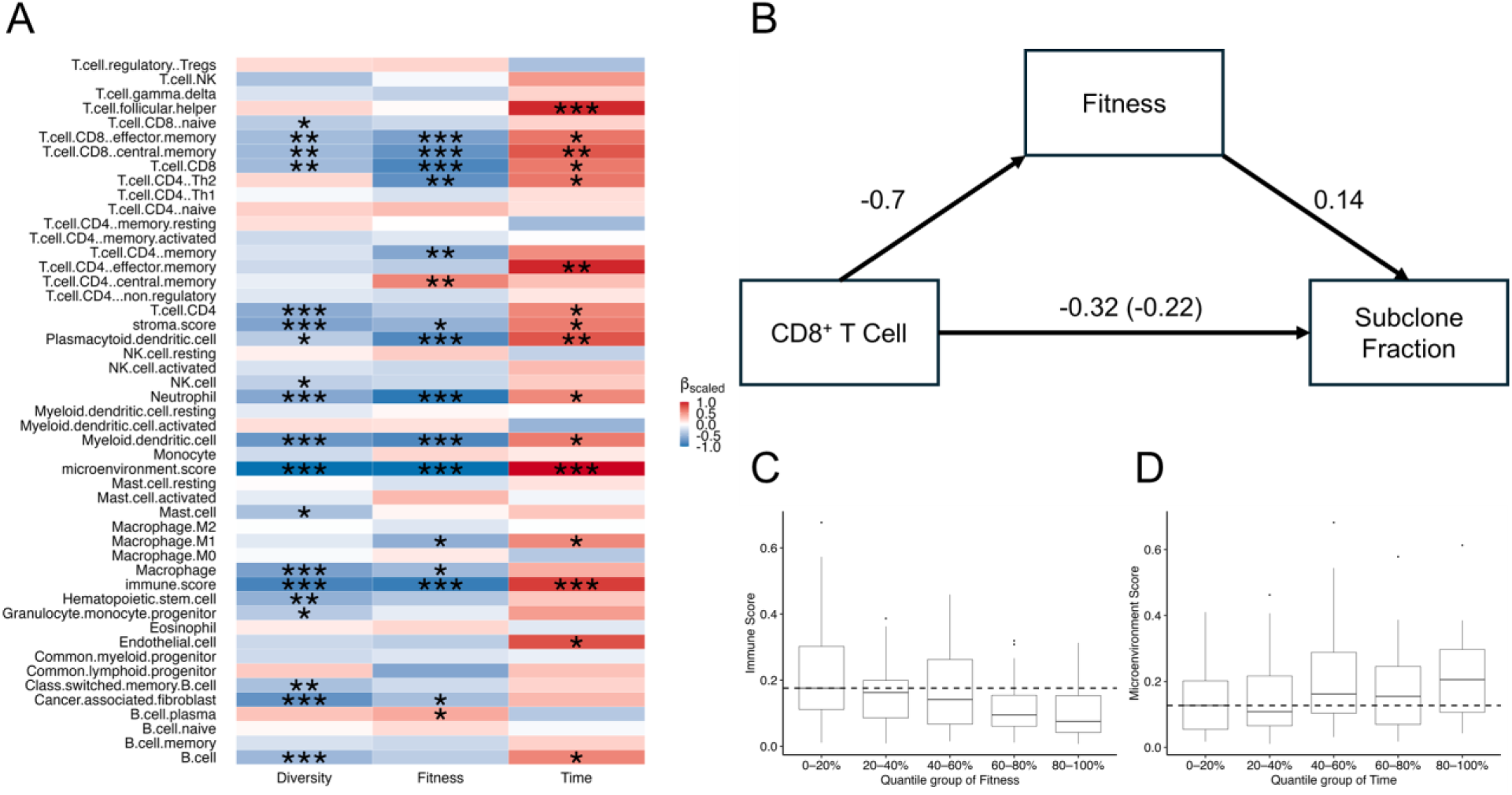
Association between evolutionary dynamics and the tumor microenvironment. (**A**) Heatmap of multivariable regression coefficients linking fitness diversity index θ, subclone fitness *s*, and subclone emergence time *t*_*f*_ to immune cell abundances; asterisks denote FDR < 0.05 (*), < 0.01 (**) or < 0.001 (***). (**B**) Mediation model with CD8⁺ T cell abundance in microenvironment as the exposure, subclone fitness as the mediator, and subclone cellular fraction as the outcome (average causal mediation effect = –0.097, *P* = 0.002; average direct effect = –0.227, *P* =0.042; total effect = –0.325, *P* =0.002; proportion mediated = 30.5%, *P* =0.004). (**C**) In LUSC, tumors with immune-hot microenvironment, as indicated by high immune score (C) and high microenvironment score (D), tend to harbor subclones with low fitness and late emergence time.

We next sought to establish the causal relationship among these parameters via mediation analysis, in which microenvironment factors served as exposures, evolutionary parameters are mediators, and subclone fraction as the outcome. The results showed that 20 immune-cell subsets significantly influenced the subclone fitness and emergence time, which in turn affected the subclone fraction (**Supplementary Table 2**). For example, CD8⁺ T cells were found to suppress subclone expansion, with 30.5% of this effect mediated through their negative impact on subclone fitness (*P* = 0.004; Fig. 5B). Tumor-type specific models revealed the same patterns (**Supplementary Table 3,** Fig. 5C**-D**), reinforcing the conclusion that immune surveillance imposes a universal constraint on subclonal outgrowth.

### Characteristics of root and branch mutations

In the TEATIME model, *MRCA*_0_ acquires growth advantage over normal cells, and *MRCA*_*f*_ gains a further advantage over the ancestral clone. If this model holds true, we would expect root mutations in *MRCA*_0_ and branch mutations in *MRCA*_*f*_ to be enriched with drivers. To test this hypothesis, we used the dNdScv package^41^ to computed dN/dS ratios for root, branch, and incidental mutations aggregated over all tumors of the same type. The background distribution was generated across 100 iterations, each randomly sampling one incidental mutation from a tumor, pooling them over all tumors, and calculating the dN/dS ratio (**Methods & Materials**). In multiple cancer types, dN/dS ratios of root and branch mutations were significantly elevated compared to incidental mutations, indicating evidence of positive selection (**Supplementary** Fig. 7). Using COAD as the example, branch mutations exhibited the highest dN/dS ratio (1.15), followed by root mutations (1.12), both of which were significantly higher than incidental mutations (1.0; t-test *P* =1.56×10^-37^ and 8.08×10^-33^, respectively, Fig. 6A). To account for the influence of gene-specific mutation rates on dN/dS ratio, we further calculated the cancer effect size of root, branch, and incidental mutations in each cancer type using the cancereffectsizeR package^5^. Consistent with the dN/dS analysis, effect size estimates followed the same pattern: branch and root mutations exhibited significantly larger effect sizes than Incidental mutations (log-scale effect size = 3.62, 3.42 and 2.77, respectively, *P* <0.05, Fig. 6B**, Supplementary** Fig. 8). We observed similar trends in BLCA, LUAD, LIHC, BRCA, HNSC, KIRC and CESC tumors (**Supplementary** Fig. 7-8).

**Figure 6.**
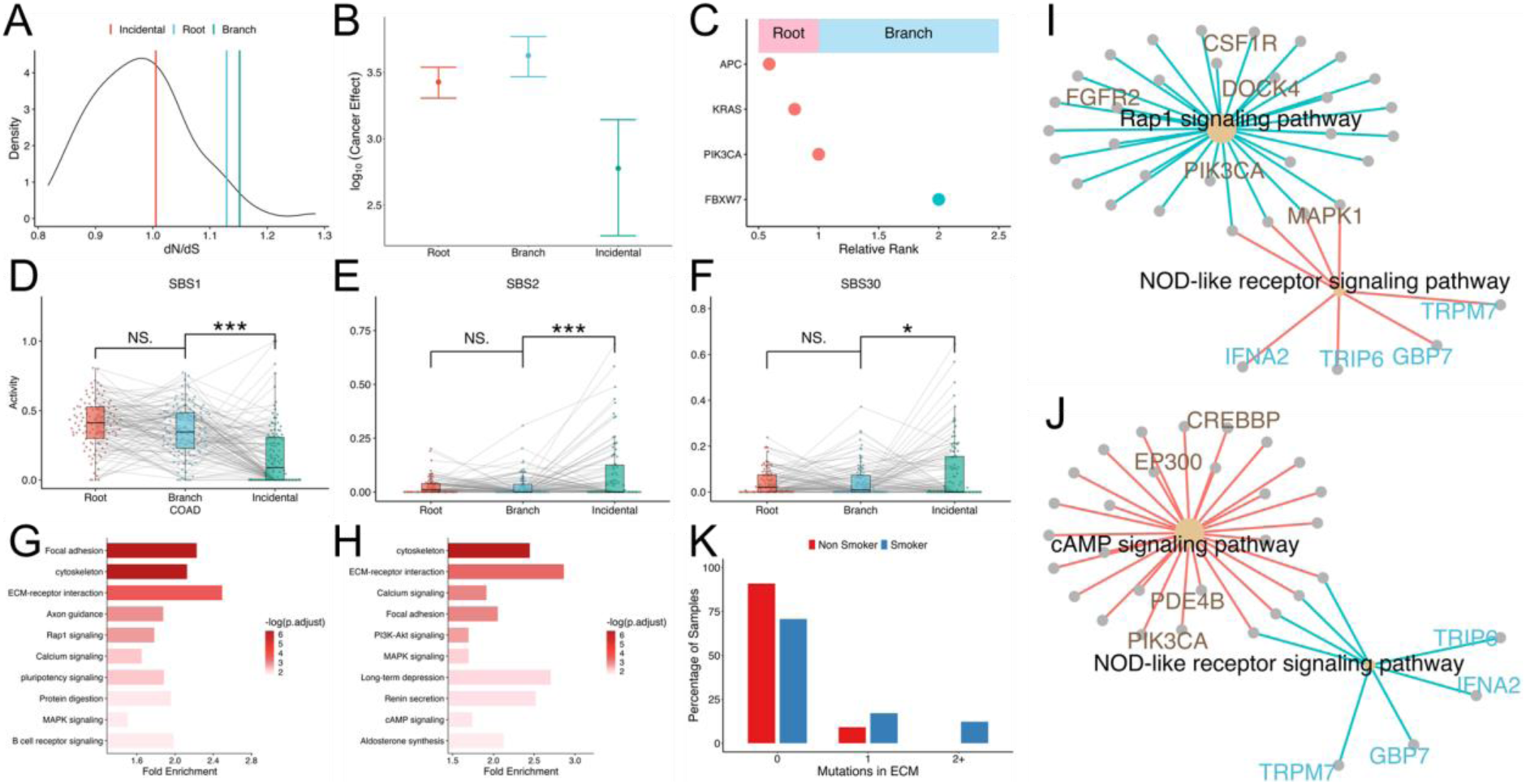
Characteristics of mutations acquired at various time points. (**A**) dN/dS ratios for root, branch, and incidental mutation; the density curve represents background dN/dS ratios calculated from randomly sampled incidental mutations. (**B**) Cancer effect size with 95% confidence interval for root, branch and incidental mutations. (**C**) Genes ranked by the ratio of their mutation frequencies in root versus branch categories. (**D-F**) Mutational signature decomposition for root, branch and incidental mutations, illustrating distinct signature profiles across temporal categories. (**G-H**) Top ten signaling pathways enriched among root-mutated genes (G) and branch-mutated genes (H). (**I-J**) Pathway network demonstrating that root mutations in key pathways increase the likelihood of downstream branch mutations. (**K**) Correlation between patient smoking status and root-mutation burden in extracellular matrix (ECM) related pathways (*P* < 0.05).

We next compared mutational signatures of root, branch, and incidental mutations. Using COAD as an example, we identified three mutational signatures with distinct activity profiles across these groups (Fig. 6D**-F**). The first signature SBS1 showed high activity in both root and branch mutations but declined significantly in incidental mutations (FDR= 1.12×10^-7^). In contrast, SBS2 and SBS30 were more active in incidental mutations than in root and branch mutations (FDR= 0.003 and 0.07, respectively). We further examined the known etiologies of these signatures. SBS1 is a clock-like signature associated with the spontaneous deamination of 5-methylcytosine. The high activity of this signature in root and branch mutations imply that mutation accumulations in the early stages of tumorigenesis is largely due to aging processes. However, as tumors progress and undergo genome-wide hypomethylation, the pool of 5-mC substrates available for deamination decrease^42,43^, potentially explaining the reduced SBS1 activity in incidental mutations. In fact, we observed a late-stage decline in SBS1 activity across five cancer types (**Supplementary** Fig. 9), consistent with the known timing of hypomethylation as a late event in these cancer types. SBS2 and SBS30 are both associated with perturbations in base excision repair, but through distinct processes^44–48^. SBS2 reflects APOBEC3 activity, while SBS30 is linked to inactivation of NTHL1. The increased activity of these signatures in incidental mutations suggests that DNA repair deficiencies become more pronounced at later stages of tumor evolution.

### Temporal relationships

To better understand the evolutionary progression leading to the emergence of the derived subclone, we examined the relationships between root and branch mutations using COAD as an illustrative example. We began by calculating the frequency of mutations in each gene classified as root or branch, and ranked genes based on the ratio of root-vs-branch frequency (Fig. 6C). This simple metric revealed that *APC* was the most frequently mutated gene in the root, followed by *KRAS* and *PIK3CA*. In contrast, *FBXW7* was most commonly mutated in the branch. This pattern aligns with the established sequence of genetic alterations in colorectal cancer progression: *APC* → *KRAS* → *PIK3CA* → *FBXW7* ^49–51^, and reinforces the role of *APC* as an early driver in colorectal carcinogenesis.

We then categorized genes into root or branch groups based on whether they were most frequently mutated in *MRCA*_0_,or *MRCA*_*f*_, respectively. Based on functional annotations from the KEGG database, we performed overrepresentation tests on root-mutated genes. The results revealed that multiple pathways were differentially perturbed at this evolutionary stage (Fig. 6G**, Supplementary** Fig. 10A). In COAD, we observed significant enrichment in canonical cancer-related pathways, including PI3K–AKT signaling and MAPK signaling, as well as several pathways related to the tumor microenvironment, such as extracellular matrix (ECM) organization and focal adhesion. Interestingly, branch-mutated genes were also enriched in a largely overlapping set of pathways (Fig. 6H, **Supplementary** Fig. 10B), suggesting that both early and late drivers tend to converge on core oncogenic and microenvironmental processes.

In contrast, incidental mutations showed no significant enrichment in any pathway, further supporting their presumed lack of functional relevance **(Supplementary** Fig. 10C).

Next, we explored temporal dependencies between root and branch mutations. For each tumor, we annotated mutations based on the functional pathways of their associated genes. We then performed linear regression to assess whether root mutations predispose specific pathways to be mutated in the branch. In CESC, we found that mutations in Rap1 signaling and cAMP signaling pathways in *MRCA*_0_ increased the likelihood of the NOD-like receptor signaling pathway being mutated in *MRCA*_*f*_. Such influence might be due to connections between these pathways via shared genes (Fig. 6I**-J**). In LUAD, mutations in ECM–receptor interaction pathway in *MRCA*_0_predisposed to mutations in the NOD-like receptor signaling pathways in *MRCA*_*f*_. Although these two pathways do not share genes, they are both involved in tumor– microenvironment interactions, suggesting functional interplay ^52–55^ ^56,57^. Notably, we found that smokers exhibited significantly more ECM-related mutations compared to non-smokers (*P* =0.047, Fig. 6K), suggesting smoking may act as an upstream factor, reshaping the tumor microenvironment to which evolving cancer clones adapt.

### Association with clinical phenotypes

Lastly, we found that evolutionary parameters have prognostic power in many cancer types. In COAD, elevated subclone fitness predicted worse prognosis (FDR =0.1, Fig. 7B). Several other cancer types also showed trends indicative of prognostic relevance at nominal *P* <0.05, although FDR corrected for multiple comparison did not reach statistical significance. In LGG, higher mutation rate was linked to poorer overall survival (Cox regression *P* =0.047, Fig. 7A). In UCEC, earlier emergence of derived subclone conferred adverse outcome (Cox regression *P* =0.013, Fig. 7C). In GBM, increased subclone expansion score τ was associated with reduced survival (Cox regression *P* =0.042, Fig. 7D). These results demonstrate that various aspects of tumor evolutionary dynamics carry prognostic information across cancer types.

**Figure 7.**
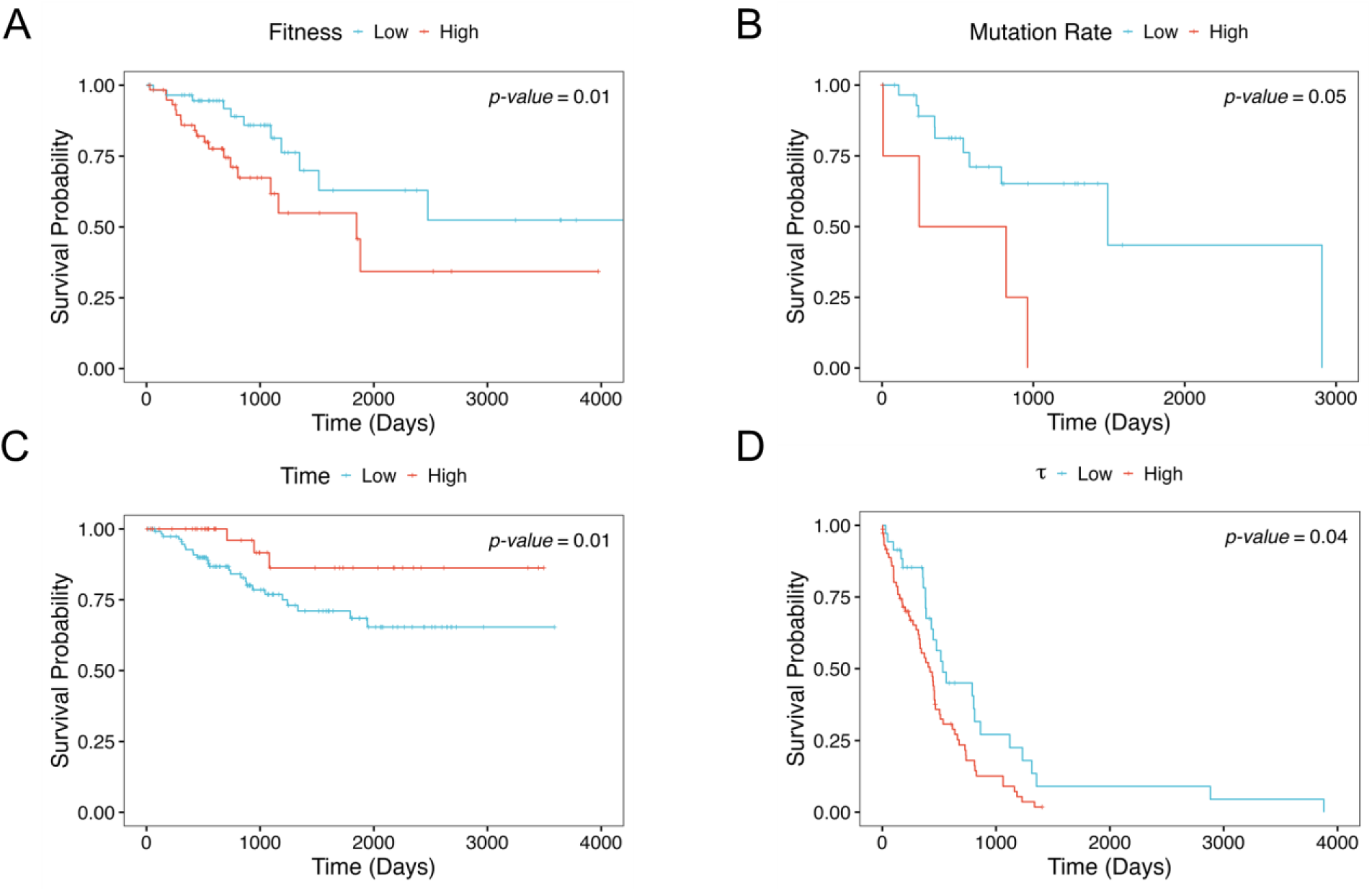
Association of evolutionary parameters with clinical features. Kaplan–Meier plots compare survival rates between two tumor groups based on various evolutionary parameters. The thresholds used to stratify tumors were determined based on the best *P* values. (**A**) COAD tumors stratified into low vs. high fitness groups. (**B**) groups LGG tumors stratified into low vs. high mutation rate groups. (**C**) UCEC tumors stratified into low vs. high subclone emergence time. (**D**) GBM tumors stratified into high vs. low τ groups.

## DISCUSSION

Understanding the evolutionary dynamics of tumors is essential for uncovering the mechanisms that drive tumor heterogeneity, progression, and therapeutic resistance. In this study, we present TEATIME, a novel computational framework that infers mutation rate, subclone fitness, and subclone emergence time from bulk sequencing data. In benchmarking analyses, TEATIME consistently outperformed existing tools in both accuracy and robustness across a wide range of tumor sizes and sequencing depths. Importantly, TEATIME yielded 2 – 37 folds of increase on the number of analyzable tumors across various cancer types, underscoring its utility for real-world clinical genomics applications.

By introducing fitness diversity as a novel measure of intratumor heterogeneity, we uncovered a universal constraint imposed by immune-inflamed microenvironments: Tumors with greater immune infiltration across multiple effector and regulatory cell types exhibit less fitness diversity. Mediation analyses reveal that this constraint operates indirectly, through reductions in subclone fitness and delayed emergence. These results integrate and extend previous reports, demonstrating that immune surveillance imposes selective pressure on tumor evolutionary trajectories by modulating both the fitness landscape and temporal ordering of emerging clones.

By comparing root, branch and incidental mutations we uncover a temporal inflection in subclone emergence. Branch mutations are under intensified positive selection and enriched in cell adhesion, ECM remodeling, and inflammatory signaling pathways. Mutational signature analysis shows that SBS2 activity increases after the onset of the more fit derived subclones while the clock-like SBS1 signature declines. These coordinated changes in selection pressure, pathway engagement, and mutational signatures define the consequences of the emergence of subclone outgrowth. These results are consistent with the Big Bang model of tumorigenesis which posits that most driver mutations occur early followed by rapid clonal expansion and later genomic stability.

Importantly, the evolutionary parameters inferred by TEATIME carry independent prognostic value. In multivariate survival models encompassing diverse cancer types, higher subclone fitness and earlier emergence both predict poorer overall survival. Thus, TEATIME not only reconstructs past evolutionary trajectories but also forecasts clinical outcomes.

TEATIME is built on assumptions of diploidy and an infinite-sites model, and it presumes comprehensive sequencing of high-frequency mutations acquired in the early foundation lineage. Violation of these assumptions will reduce the accuracy of the estimations. It currently models cancer cell growth as a clonal evolution model with exponential growth. In tumors that deviate from these models, such as those follow tumor stem cell model and those with low growth rate in the core of a tumor than proliferative rims, or those with logistic growth due to resource constraints, TEATIME will not converge or produce biased estimates. Beyond these modeling constraints, all tools require a sufficiently large number of mutations to ensure robust performance. Therefore, we do not recommend using these tools to analyze tumors with low mutation burden or those assessed using targeted sequencing. In fact, TEATIME failed to converge for approximately 80% of TCGA tumors due to these constraints. We further note that MOBSTER was developed for WGS, which may limit its performance on WES datasets.

Additionally, because bulk sequenced biopsies capture only a limited region of the tumor, the identified subclones may reflect spatially localized populations rather than the complete evolutionary landscape. Extending the framework to model multiple coexisting subclones, spatial heterogeneity, and non-exponential growth dynamics, as well as incorporating longitudinal or single cell sequencing data, will broaden its applicability.

## Acknowledgments

We thank Trevor Graham, Calum Gabbutt, and Giulio Caravagna for their helpful discussions and valuable insights throughout this work. This work was supported by NIH R01LM013438.

## METHODS AND MATERIALS

### Cell proliferation parameters

The parameter λ controls the cell birth rate. By setting λ = ln 2, it indicates that the division of a parent cell produces two daughter cells. The parameter β = 1 + ^ln Φ^⁄ λ, where Φ defined as cell birth rate – cell death rate. Users can specify the value of β. By default, we set β = 0.9.

### Input data and preprocessing

The input data for TEATIME has three columns: the read count of the wildtype allele, the read count of the mutant allele, and the ploidy of the genomic region harboring the mutation. Only heterozygous mutations located in diploid regions are used. To estimate the sample impurity, VAFs are clustered using the MAGOS program and the mean VAF is calculated for each cluster. The highest mean VAF Φ provides an estimate of the sample purity ρ = 2Φ. To adjust for sample impurity, the VAF of each mutation is adjusted to *VAF*(2 − ρ)/(2*VAF*(1 − ρ) + ρ).

### Initial decomposition of VAFs

Mutations are first split into two partitions using the MAGOS program^58^. Based on beta-distributions reparametrized with mean sequencing depth and mean VAF, MAGOS progressively groups mutations into a hierarchical tree structure. TEATIME traverses this tree, calculating the mean VAF for each node. The first partition Ω_1_ aggregates mutations in nodes with mean VAF>0.25, indicating presence in >50% of cells. The second partition Ω_2_contains the remaining mutations.

Within each partition, mutation are further split into clusters using the automixfit() function from the R/RBesT package^59^. The automixfit() function is configured with Nc=seq (1, 10) and k=6, modeling the VAFs as a mixture of beta distributions with 1 to 10 components. The shape parameters α and β for each identified component are returned. Using *v* = β/(α + β), each component is reparametrized as ℬ(*v*). Mutations are then assigned to one of these components, which satisfies argmax_*x*∈*v*_ *prob*(*VAF*|ℬ(*v* = *x*)), where *v* is the collection of *v* values of the identified components. Mutations assigned to the same component form a cluster, with a mean VAF *v*^-^. It is noteworthy that while the *v*^-^ value of a cluster is close to the *v* value of the corresponding component, they are not necessarily equal. Furthermore, automixfit() requires the number of components is prespecified, which is unknown. Therefore, these clustering results provide only initial starting points for further optimization.

### Joint optimization of mutation clusters and evolutionary parameters

Given the large number of unknowns, including mutation clusters and evolutionary parameters, an exhaustive search for optimal solutions is intractable. To address this problem, TEATIME carefully defines the search spaces for a subset of variables, iteratively selects possible values from these spaces, and uses them to estimate the remaining variables. Solutions from each iteration are compared, and the one showing the best fit between the observed VAFs and the inferred evolutionary process is selected as the final solution.

Specifically, if the branch cluster *U*^*b*^∼ℬ(*v*^*b*^) is located, *P* can be estimated as 2*v*^*b*^. Given the *P* value, *v*^*k*^ can be calculated for each *t* ∈ [1,2, …, *t*_*f*_ − 1] and applying *f*_*k*_(*t*) = 2*v*^*k*^. Based on the series of *v*^*k*^ values, mutations can be grouped into a set of trunk clusters *U*^*k*^∼ℬ(*v*^*k*^). The μ value can then be inferred either by counting mutations in the *U*^*k*^ cluster or by estimating the slope of a line fitted across all *U*^*k*^ clusters based on Eq. (4).

We first define the search space for *v*^*b*^. In a tumor dominated by *K*_*f*_, i.e., *P* > 0.5, it is expected that *v*^*b*^ > 0.25, and the *U*^*b*^ cluster is in the Ω_1_ partition. Among all clusters in Ω_1_, Root cluster *U*^*r*^ cannot be *U*^*b*^. To find *U*^*r*^, the cluster with the highest mean VAF *v*^-h-^ is examined. If *v*^-h-^∈ [0.45, 0.55], this cluster is assigned as *U*^*r*^. Otherwise, a new component ℬ(*v* = 0.5) is created. Each VAF in Ω_1_ is reassigned to one of the existing and new components, which satisfies argmax_*x*∈{V,0.5}_ *prob*(*VAF*|ℬ(*v* = *x*)). The new cluster *U*∼ℬ(*v* = 0.5) is again split using the automixfit() function, and the cluster with the highest mean VAF is examined. This process continues till the *v*^-h-^ falls within the range [0.45, 0.55]. Upon completion, three clusters are recorded – *U*^*r*^ is the cluster with the highest mean VAF; *U*^′^ is the cluster with the second highest mean VAF *v*^-^′^-^; and *U*^′′^ the one with lowest mean VAF *v*^-^′^-^′^-^. Theoretically, *v*^*b*^ can be any value in the range [*v*^-^′^-^, ^-^*v*^-^′^-^′^-^]. Practically, to reduce computation load, only *v*^-^′^-^, *v*^-^′^-^, and (*v*^-^′^-^ + *v*^-^′^-^′^-^)/2 are considered.

In a tumor dominated by *K*_*a*_, it is expected that *v*^*b*^ < 0.25, and *U*^*b*^ is the cluster with the highest mean VAF in the Ω_2_ partition. Starting with the initial clusters in Ω_2_, the one with the highest mean VAF is iteratively split using the automixfit() function until no new clusters can be produced. Upon complete, and the cluster with the highest mean VAF *v*^-^′^-^′^-^′^-^ is assigned as *U*^*b*^.

The above steps produce four possible *v*^*b*^ values: *v*^-^′^-^, *v*^-^′^-^, (*v*^-^′^-^ + *v*^-^′^-^′^-^)/2, and *v*^-^′^-^′^-^′^-^, which constitute the search space for *v*^*b*^. The search space for *t*_*f*_ includes all integers between 1 and the total number of mutations in Ω_1_ that are outside the *U*^*r*^ cluster (i.e., non-root mutations).

For a given pair of *v*^*b*^ and *t*_*f*_ values selected from their respective search space, *P* and a series of *v*^*k*^ values, denoted as *v*^*k*^, are estimated (Supplementary Materials).Non-root mutations are grouped into clusters, where the VAF of each mutation satisfies argmax _*b* *k*_ *prob*(*VAF*|ℬ(*v* = *x*)). Based on the identified *U*^*k*^ clusters, two approaches are used to estimate μ.

The mutation counting approach relies on the *U*^*k*^ cluster, because this cluster has the highest mean VAF among all *U*^*k*^ clusters, making it less likely to be mixed with incidental mutations and more likely to be exhaustively sequenced (Supplementary Materials). Given the *U*^*k*^ cluster with a mean VAF *v*^-^*k*^-^, *k* observed VAFs closet to *v*^-^*k*^-^ are selected. Meanwhile, *k* simulated VAFs are randomly sampled from the ℬ(*v* = *v*^-^*k*^-^) distribution. If the *k* observed VAFs come from trunk mutations acquired during the same cell division, their distribution should be similar to that of the *k* simulated VAFs. This hypothesis is tested using the Mann-Whitney test with a type-I error cutoff of 0.05 ^60,61^. A series of *k* values are evaluated, ranging from 1 to the number of mutations in *U*^*k*^. Those accepting the null hypothesis are retained as possible estimates for μ. If more than one *k* value is retained, they are ranked by the effect size measured by Cliff’s delta score.

The line-fitting approach uses all *U*^*k*^ clusters. Based on Eq. (4), a line is fit between − ln(2*v*^-^*k*^-^ − *P*) and *M*_*k*_(*t*), where *v*^-^*k*^-^ is the mean VAF of cluster *U*^*k*^ and *M*_*k*_(*t*) is cumulative counts of trunk mutations till time *t*. The mutation rate is calculated as μ = λβγ, where γ is the slope of the fitted line. Meanwhile, a set of simulated clusters are created by randomly sampling μ VAFs from each ℬ(*v* = *v*^-^*k*^-^) distribution. A line is fit across these simulated clusters and the μ^ is estimated from the slope using the same procedure described above. This simulated is repeated 1000 times, producing 1000 μ^ values. One-group t test is used to compare the 1000 μ^ values with the μ value estimated from the observed VAFs. If no significant difference is found, the μ value is retained.

The μ values estimated using the two approaches are compared. If consensus exists, the associated set of mutation clusters and evolutionary parameters are retained. If there is no consensus, line-fitting results are retained. If one approach produces an empty set, results from the other approach are retained.

Each pair of *v*^*b*^ and *t*_*f*_ values produce estimates for mutation clusters and evolutionary parameters. After all pairs of values are tested, these estimates are compared on Bayesian Information Criterion scores *BIC* = −2 log(ℒ) + *k* log(*m*), where ℒ is the likelihood that the observed VAFs belong to the set of ℬ(*v*^*k*^) distributions, *k* is the number of *U*^*k*^ clusters, and *m* is the number of mutations in *U*^*k*^ clusters. The set of estimates with the best BIC score is selected as the final solution.

Once *P* is known, *s* can be calculated using the mean VAF of cluster *U*^*s*^, formed by successor mutations acquired at time *t*_*f*_ + 1, based on Eq. (5). In a tumor dominated by the *K*_*f*_ subclone, the *U*^*s*^ cluster is expected to be in the Ω_2_ partition. TEATIME searches for this cluster in the initial clusters identified in Ω_2_. Given an arbitrary cluster in Ω_2_ that has a mean VAF < *v*^*b*^, it is assumed to be *U*^*s*^, and *s* is calculated (Supplementary Materials). The number of *U*^*s*^ clusters, each consisting of successor mutations acquired at a time point before *t*_*f*_ + 1, is given by *I* = *round*(*s*). Based on the series of clusters *U*^*s*^, where *i* = 1, 2, …, *I*, all mutations between *U*^*b*^ and *U*^*s*^ are regrouped to clusters. Using the updated *U*^*s*^, a new selection coefficient value *s*^′^ can be estimated. If |*s* − *s*^′^| > 10^−6^, the *s* value is updated to *s*^′^, and the steps of identifying *U*^*s*^ clusters and computing *s*^′^ are repeated. The iterations end when |*s* − *s*^′^| ≤ 10^−6^ or 1,000 iterations are completed. Lastly, *t*_*e*_ is calculated using Eq. (3).

TEATIME does not estimate *s, t*_*e*_ for tumors dominated by the *K*_*a*_ clone because the *U*^*s*^ cannot be located in the current framework.

### Simulations

The performance of TEATIME and other methods was evaluated using two sets of simulated data. The first dataset is part of the MOBSTER package^12^, consisting of 150 synthetic tumors that follow various evolutionary trajectories. While all tumors contain 10^8^ cancerous cells at the time of sampling, the other evolutionary parameters used to simulate tumor growth vary in a broad range, including *t*_*f*_ ∈ [4, 14], *s* ∈ [0.125, 1.625] and *P* ∈ [0, 0.97].Some of these tumors are monoclonal neutral, containing no subclones, while others include one subclone. All simulated samples consist of 100% cancerous cells and are sequenced at a depth of 120×.

The second dataset was generated using TEMULATOR^16^, where we varied the sequencing depth from 60 × to 1000× and varied *N* from 10^6^ to 10^7^, and *t*_*f*_ ∈ [4, 14], *s* ∈ [0.125, 1.625] and *P* ∈ [0, 0.92]. The purity is 100% for simplicity. For each set of parameter values, 150 synthetic tumors were simulated. A total of 2,100 tumor samples were simulated. Each sample contains at least 1000 mutations.

### TCGA data

We downloaded the mc3.v0.2.8.PUBLIC.maf.gz file from the Genomic Data Commons data portal^17^, which contained somatic mutations in 10,295 tumors of 33 cancer types from TCGA PanCanAtlas mapped to the human genome build hg37. After removing hyper- and and hypo-mutated tumors, defined as the those with the total number of mutations outside the 1.5 interquartile range^62^, and removing relapsed and metastatic tumors, 8,935 primary tumors of 33 cancer types were retained. For each tumor, clinical information, CNVs, and annotated somatic mutations from WES were downloaded. To ensure sufficient statistical power, we restricted downstream analyses to cancer types with at least 20 predictable samples, resulting in 19 tumor types.

### Quantification of intra-tumor heterogeneity

To quantify intra-tumor heterogeneity, we computed the fitness diversity index (θ) as the Shannon entropy of the cell population distribution. Let *P* denote the proportion of K_*f*_ cells, we then calculated

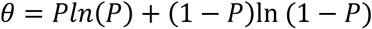

### Statistical tests of associations

Cox proportional hazards model was used to test the association between evolutionary parameters and patient overall survival. Covariates included age, gender, tumor stage, mutation load, presence of nonsynonymous mutations in driver genes if applicable. For each cancer type, driver genes were identified using the GUST program^62^, not annotated as passenger genes in the Cancer Gene Consensus^63^. Presence of nonsynonymous mutations in oncogenes in the corresponding cancer type was coded as 1 and 0 otherwise. The coxph() function in the R/survival package was used for this analysis. For Kaplan–Meier visualization, tumor stratification thresholds were defined using optimal cut points identified by cut_survpoint().

Logistic Regression was used to test the association between evolutionary parameter and cancer type/Treatment Resistance, Covariate adjustment was performed in the same manner as in the Cox proportional hazards models. For Treatment Resistance, Progressive disease (PD) in treatment outcome was treated as a binary outcome (PD = 1, non-PD = 0).

Linear regression was used to test the association between evolutionary parameter and immune infiltration, Models for the pan-cancer cohort included age and cancer type as covariates, whereas tumor specific analyses also incorporated gender, tumor stage and mutation load if applicable. P-values were adjusted using the Benjamini–Hochberg (BH) method.

### Mediation tests in immune infiltration

In the mediation test, microenvironment factors served as exposures, evolutionary parameters are mediators, and subclone fraction as the outcome. R mediation package is used for estimating indirect effect^64^.

### Selection intensity and dN/dS ratio Calculation

To evaluate dN/dS ratio, we applied the dNdScv R package^41^. We consider three locations: Root mutations, Branch mutations, and Incidental mutations. To construct a background distribution, we picked a random incidental mutation per tumor as the seed and included additional incidental mutations with VAFs ±0.01 of that seed, aggregated mutations over all tumors, computed dN/dS ratio and repeated for 100 iterations. A one-sample t-test was performed to evaluate whether the Root, Branch dN/dS ratios were significantly higher than the background dN/dS, estimated from randomly selected Incidental mutation. For Root mutations, Branch mutations, and Incidental mutations, we also calculated cancer effect size using the cancereffectsizeR package^5^.

### Mutational signature extraction and statistical test

Following mutational signature analysis practices^65^, we extract mutational signatures at root, branch and incidental evolutionary stages by excluded SBS signatures considered biologically implausible or irrelevant for each cancer type^66,67^. We then applied the MutationalPatterns package to extract mutational signatures at each location using a curated subset of cancer type–specific SBS signatures^68^.

After signature extraction, we performed paired t-tests to compare the relative contribution of each SBS signature between root and branch, and between branch and incidental mutations. P-values were adjusted using the Benjamini–Hochberg (BH) method.

### Temporal order of mutated gene

For a gene with recurrent nonsynonymous mutations (observed in at least two samples at both root and branch locations), we defined 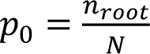 and 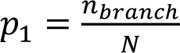, where *n_root_* and *n_branch_* are the numbers of mutations at the root and branch locations, respectively, and *N* is numbers of all mutations. We calculated the ratio ^*p*1^/_*p*_ to assess the likelihood of mutation timing.

### Temporal pathway enrichment and association

In pathway enrichment analysis, we assigned each gene to Root, Branch or Incidental if at least 50% of its nonsynonymous mutations occurred at that location and retained only those genes for each location. The remaining genes were subjected to KEGG pathway enrichment analysis using clusterProfiler^69,70^.

In pathway association analysis, we focused on nonsynonymous mutations in two locations— Root mutations and Branch mutations. Using curated KEGG pathway definitions, we constructed all possible pathway pairs. For each pair, we quantified the number of mutations occurring at Root and Branch. Pathways mutated in less than 1% of samples were excluded to focus the analysis on recurrently altered pathways. We then modeled the number of mutations in each pathway at t₁ as the response variable and those at t₀ as predictors. For pathway pairs that have shared genes, P-values were adjusted using the Benjamini–Hochberg (BH) method. For pathway pairs without shared genes, we relaxed the significance threshold by using unadjusted p-values and selected one pair as the representative. Temporal dependencies were interpreted based on the direction and magnitude of regression coefficients: positive slopes indicated synergistic relationships, while negative slopes suggested antagonism. For clarity, we focused on synergistic relationships, defined as pathway pairs with positive regression coefficients.

### Execution of programs

We downloaded the MOBSTER program from https://github.com/caravagnalab/MOBSTER. When applying MOBSTER to a tumor, the auto_setup parameters were set to FAST. We downloaded the TumE program from https://github.com/tomouellette/TumE. When applying TumE to a tumor, the parameters were set to default.

## Code availability

The R implementation of the TEATIME algorithm is publicly available on Github https://github.com/liliulab/TEATIME.

## Data availability

Only publicly available data were used in this study, and data sources and handling of these data are described above.

