## SUPPLEMENTARY_MATERIALS for "Competing Subclones and Fitness Diversity Shape Tumor Evolution Across Cancer Types"

**Approximation of tumor cell population size**: At the time point $t_e$ when a tumor is sampled for analysis, the total number of cells $N = \mathrm{N}_a + \mathrm{N}_f - N_l$ where $\mathrm{N}_a$ and $\mathrm{N}_f$ are the population sizes of $\mathrm{K}_a$ and $\mathrm{K}_f$, respectively, and $N_l$ is the number of cells that are counted twice when computing $\mathrm{N}_a$ and $\mathrm{N}_f$ and needs to be subtracted. By setting $t_0 = 0$, the tumor cell population size at $t_e$ is

$$N = \mathrm{N}_a + \mathrm{N}_f - N_l = e^{\lambda\beta t_{end}} + e^{\lambda\beta(1+s)(t_e - t_f)} - e^{\lambda\beta(t_e - t_f)}$$

When the final tumor cell population size $N = \mathrm{N}_b + \mathrm{N}_f - N_l$ is approximated as $\widetilde{N} = \mathrm{N}_b + \mathrm{N}_f$, it leads to overestimation. Here we quantify the error as

$$error = \frac{N_l}{N} = \frac{1}{e^{\lambda\beta t_f} + e^{\lambda\beta s(t_e - t_f)} - 1}$$

Let $\lambda = \ln 2$ and $\beta = 1$. When $t_f > 5$ or $s(t_e - t_f) > 5$, the error is less than 1/31. Given that tumor cells undergo a large number of cell division cycles, it is reasonable to assume the error is negligible.

**Estimation of Evolutionary parameters in root, trunk and successive clusters:** Assuming a tumor sample contains solely cancerous cells after adjusting for normal cell contamination, TEATIME models VAFs as a mixture of beta distributions. Each component in the mixture is a beta distribution $\mathcal{B}(\alpha, \beta)$ with the shape parameters $\alpha = dv$ and $\beta = d(1 - v)$, where $d$ is the mean sequencing depth and $v$ is the mean VAF [15]. Since mutations in a tumor share similar, known sequencing depths, the beta distribution can be reparametrized as $\mathcal{B}(v)$. TEATIME decomposes the mixture to identify clusters of mutations that belong to different categories. We split VAF into two partitions: first partition $\Omega_1$ aggregates mutations with VAF>0.25, indicating presence in >50% of cells. The second partition $\Omega_2$ contains the remaining mutations. The observed VAF distribution in $\Omega_1$ is modeled as a three component Beta-mixture:

$$p(VAF) = \pi_r \mathcal{B}(v^r) + \pi_t \sum_{t=1}^{t1-1} \mathcal{B}(v_t^k) + \pi_b \mathcal{B}(v^b)$$

subject to

$$\pi_r + \sum_{t=1}^{t1-1} \pi_t + \pi_b = 1$$

Where Root mutations form a cluster $U^r \sim \mathcal{B}(v^r)$. Trunk mutations, acquired at various time points, generate a series of clusters $U_t^k \sim \mathcal{B}(v_t^k)$ ,where $f_k(t) = 2v_t^k = P + (1-P)\frac{1}{e^{\lambda\beta t}}$. Branch mutations, representing later evolutionary events, form a single cluster $U^b \sim \mathcal{B}(v^b)$. Here, $\{\pi_r, \pi_t, \pi_b\}$are the relative contributions and are inferred by maximum likelihood from the data. The fraction of $K_f$ cells among all tumor cells is estimated as $f_h = 2v^b$.

The mutation rate is inferred from the trunk clusters using two approaches. The first approach quantifying the number of mutations in the earliest trunk cluster $U_{t=1}^k$. Assuming all mutations in this cluster have been sequenced, the number of variants gives an estimate of μ. The second approach aggregates information over multiple trunk clusters. Based on Eq. (4), a line can be fitted between $-\ln(f_k(t) - P)$ and $M_k(t)$ across the series of $U^k$ clusters. The slope of this line is given by $\gamma = \mu/\lambda\beta$, where $\lambda = \ln 2$ and β is a user-specified cell survival rate between 0 and 1. Consensus between two approaches is taken as the final estimates. Based on Eq. (4), $t_f = M_k/\mu$.

The selection coefficient $s$ is inferred using successive mutations in $\Omega_2$. Because the expected VAFs of successive mutations acquired at $t > t_f + 1$ are lower than $P/4$, and clusters formed by low-frequency mutations are difficult to identify, we limit the analysis to successive mutations acquired before or at $t_f + 1$. Compared to $K_a$ cells that go through one generation of cell division in a unit time, the fitness advantage of $K_f$ cells is reflected in the $MCRA_f$ cell and its descendants undergoing $1 + s$ generations of cell division between time points $t_f$ and $t_f + 1$.

Each generation produces a cluster, including the cluster $U^s_{t_{f+1}}$ that corresponds to mutations acquired at $t_f + 1$ and additional $I = round(s)$ clusters that correspond to mutations acquired in intermediate generations before $t_f + 1$. Based on Eq. (5), the selection coefficient can be estimated as

$$s = \frac{\ln\left(\frac{P}{2v^s_{t_f+1}}\right)}{\lambda\beta}$$

Subsequently, $t_e$ can be calculated using Eq. (3).

The above estimations assume that the $U^r$, $U^k$, $U^b$, and $U^s$ clusters have already been identified. However, in practice, locating these clusters is challenging. TEATIME employs a joint optimization algorithm that identifies informative clusters and estimates evolutionary parameters concurrently, maximizing the likelihood that the observed VAF distribution is generated by the inferred evolutionary process (**Supplementary Fig. 1, Methods and Materials**).

**Mixing of incidental mutations in root and trunk clusters**: If $K_f$ is dominant, all mutations from the foundation lineage will reside in the $\Omega_1$ partition. Some incidental mutations acquired by early progeny of the $MRCA_0$ cell and some successor mutations acquired by early progeny of the $MRCA_f$ cells may also fall into the $\Omega_1$ partition due to the death of sister cells.

Here we quantify the extent of mixing of incidental mutations in the $U^k_{t=1}$ and $U^r$ clusters. Let $X$ represent the VAFs of incidental mutations generated by the $i$-th cell division in $\mathrm{K}_a$, with the mean VAF as $f_i$. Let $Y$ represents the VAFs of trunk mutations acquired at time $t = 1$, with the mean VAF as $f_1$. If cell proliferation rate is 2, $f_1$ is always higher than $f_i$ , i.e., $\Delta_\mathrm{i} = f_i - f_1 < 0$, as these trunk mutations are present in all $\mathrm{K}_f$ cells and half of $\mathrm{K}_a$ cells. The probability that an incidental mutation acquired at $i$-th cell division mixes into $U^k_{t=1}$ and $U^r$ is $prob(X > Y)$. To get a close-form approximation of this probability, a normal distribution approximation is employed[1] as

$$prob(X > Y) \approx \Phi\left(\frac{\Delta_i}{\sqrt{\sigma_x^2 + \sigma_y^2}}\right)$$

where $\Phi$ (.) is the cumulative density function of normal distribution $\mathcal{N}(0, 1)$. Given that the sequencing depth is typically significantly greater than 1, the variance of the beta distribution can be approximated as $\frac{v(1-v)}{d}$. Therefore,

$$\Phi\left(\frac{\Delta_i}{\sqrt{\sigma_x^2 + \sigma_y^2}}\right) = \Phi\left(\frac{\Delta_i \times \sqrt{d}}{\sqrt{2f_1(1 - \Delta_i - f_1) + \Delta_i - \Delta_i^2}}\right) \leq \Phi\left(\frac{\Delta_i \times \sqrt{d}}{\sqrt{\frac{1 - \Delta_i^2}{2}}}\right)$$

Although $f_1$ cannot assume the maximum value, the above estimate offers a way to establish an upper bound for the probability of mixing. The probabilities of VAFs of incidental mutations mixing into $U_{t=1}^k$ vary with the value of $i$ and the proportion of the $K_f$ subclone $P$.

We performed simulations to assess the mixing probability for a series of $P$ values from 10% to 40% and sequencing depths from 30× to 1000×. We then estimated the $prob(X > Y)$ by taking random samples from the beta distribution (reflecting a realistic scenario) and from a normal distribution approximation (acting as a closed-form upper bound). As expected, the simulations show that the normal distribution approximation effectively provides an upper bound for estimating the level of mixing (**Supplementary Fig. 2**). When d=100 and survive rate = 1, it is straightforward to see that when $i$ is 1, the probability of mixing is 0.01595 if $P = 30\%$, and 0.23947 if $P = 10\%$. When $i$ increases to 2, these probabilities change to 0.00027 for $P = 30\%$ and 0.00993 for $P = 10\%$. As $i$ becomes sufficiently large, the probability of mixing decreases significantly, reaching as low as 5.3x$10^{-9}$ for $P = 30\%$, and 0.00013 for $P = 10\%$.

Even if the proportion of $K_f$ is sufficiently small, at most $0.25\mu$ incidental mutations will be mixed into the $U_{t=1}^k$ cluster. However, this scenario is often impractical in actual cancer samples[2], suggesting that $U_{t=1}^k$ despite the potential for minor mixing, remains the most distinct and least contaminated cluster in the $\Omega_1$ partition.

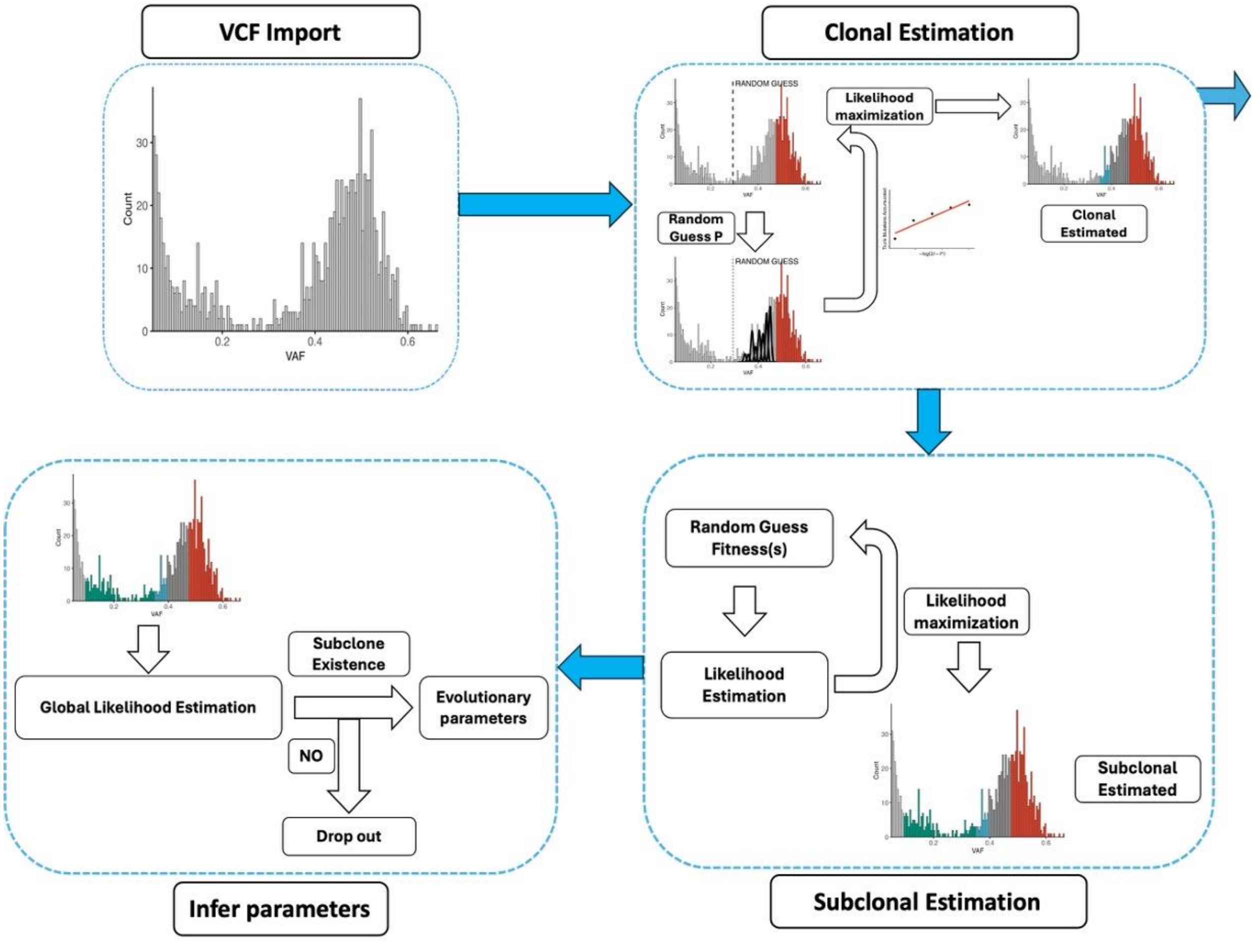


**Supplementary Figure 1.**
Flow chart describing the joint optimization process.

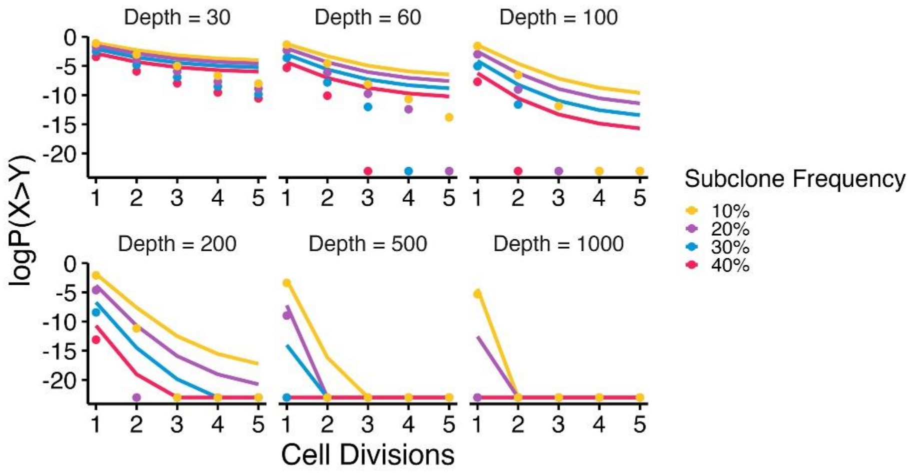


**Supplementary Figure 2**. Estimation of the probability of incidental mutations mixing into the root or first trunk clusters. The probability of mixing decreases with the cell division generations and sequencing depths. The line represents the normal distribution approximation, while the data points are derived from Beta distribution sampling. For clarity in visualization, probability $<10^{-10}$ is set to $10^{-10}$.

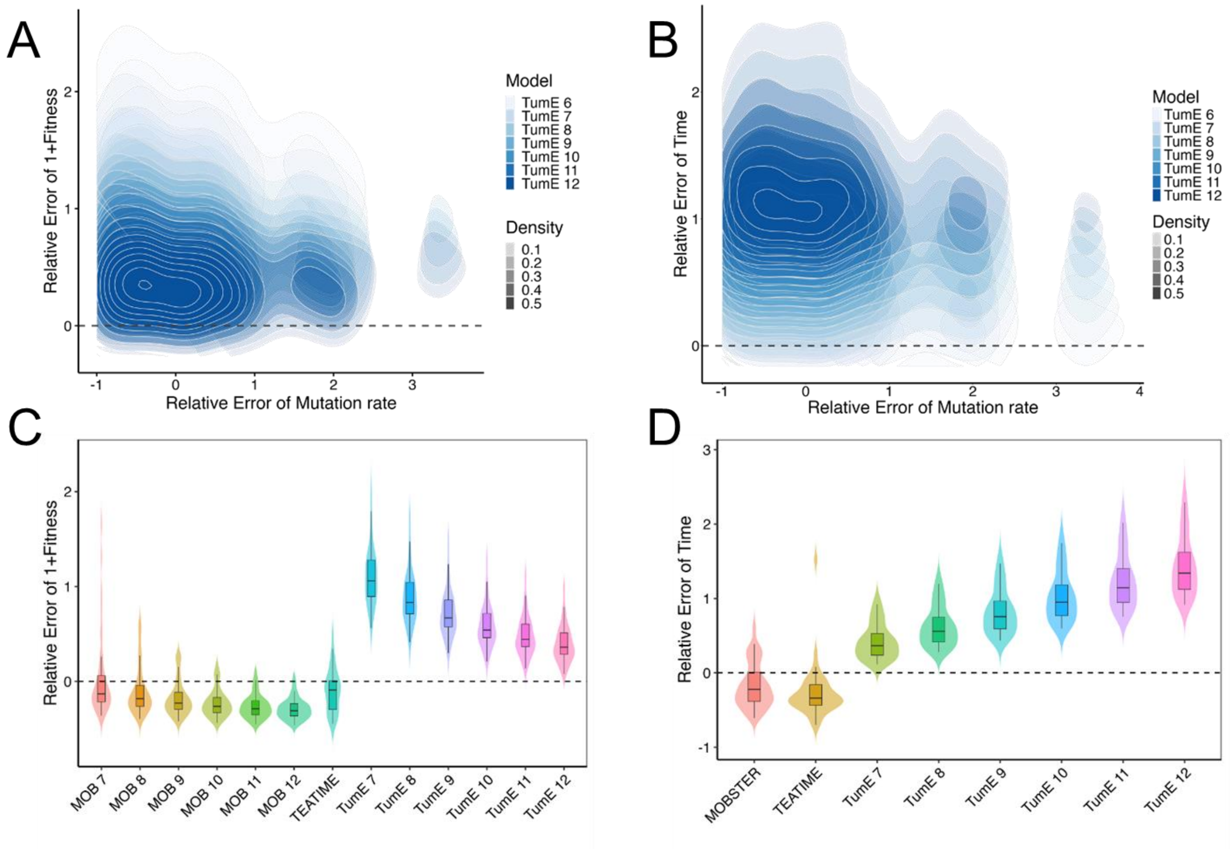


**Supplementary Figure 3.** Relative error of parameter estimates across different population sizes (N) for TumE, MOBSTER, and TEATIME.

(A-B) Relative errors of TumE Monte Carlo estimates for mutation rate (μ), fitness (s), and subclonal emergence time ($t_1$) across input population sizes ranging from $10^6$ to $10^{12}$. Colors represent different values of N.

(C-D) Comparison of relative errors in s, and $t_1$ estimates obtained from TumE (mean prediction), MOBSTER, and TEATIME. MOBSTER fitness prediction results at N = $10^6$ were excluded due to instability and extreme error values.

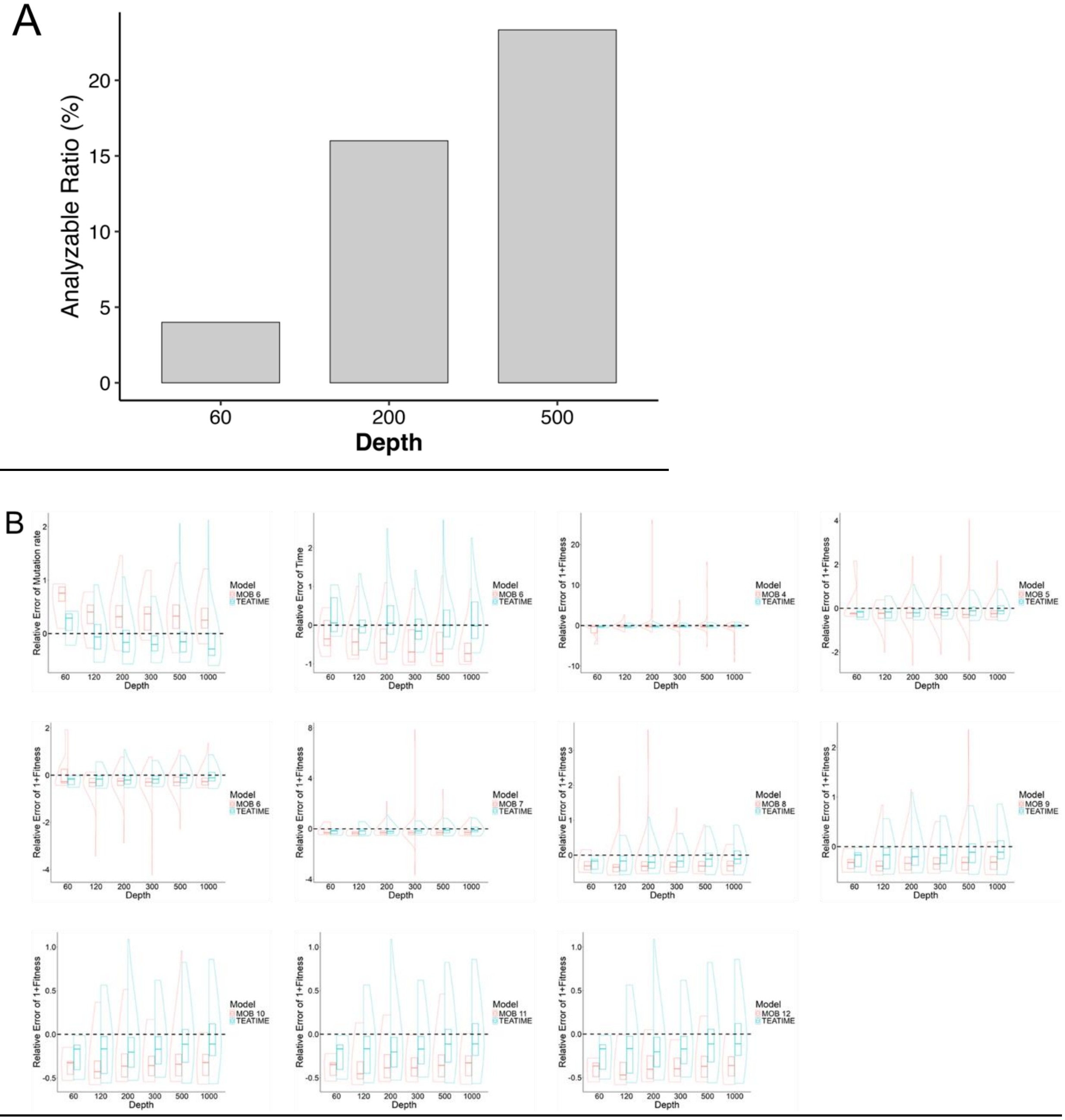
A
Analyzable Ratio (%)
0
5
10
15
20
60
200
500
Depth
B
Relative Error of Mutation rate
Relative Error of Time
Relative Error of 1+Fitness
Depth
60
120
200
300
500
1000
Model
MOB 4
MOB 5
MOB 6
MOB 7
MOB 8
MOB 9
MOB 10
MOB 11
MOB 12
TEATIME

C

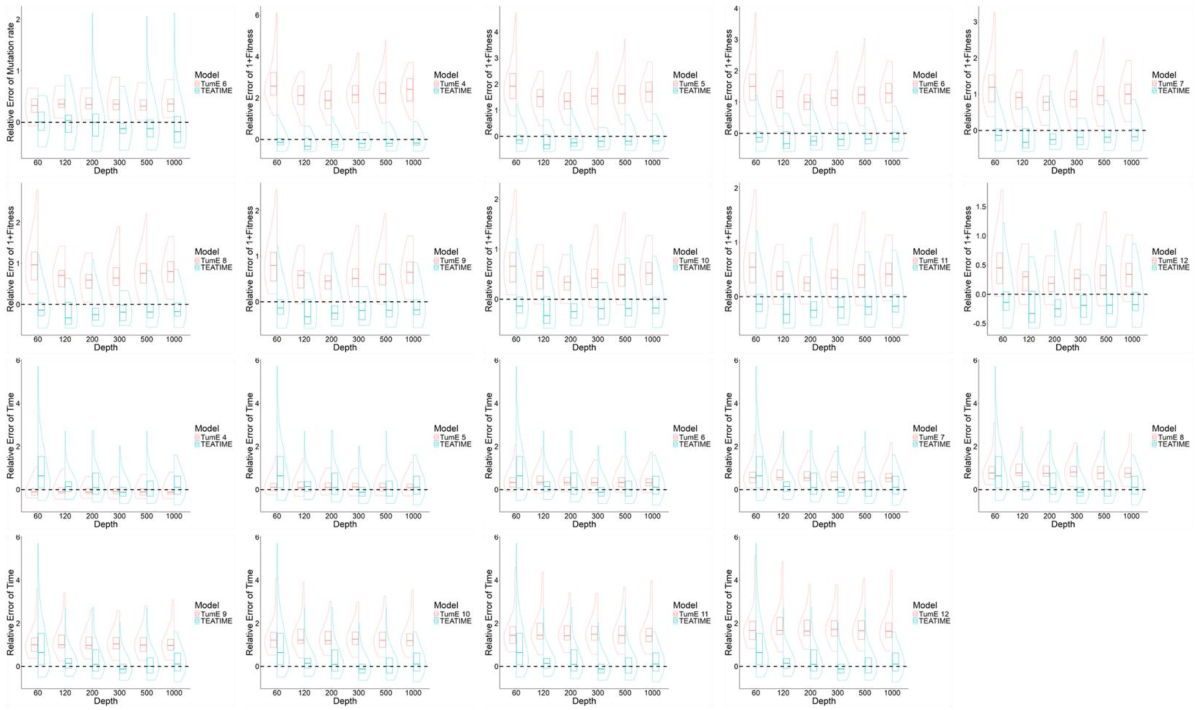

D

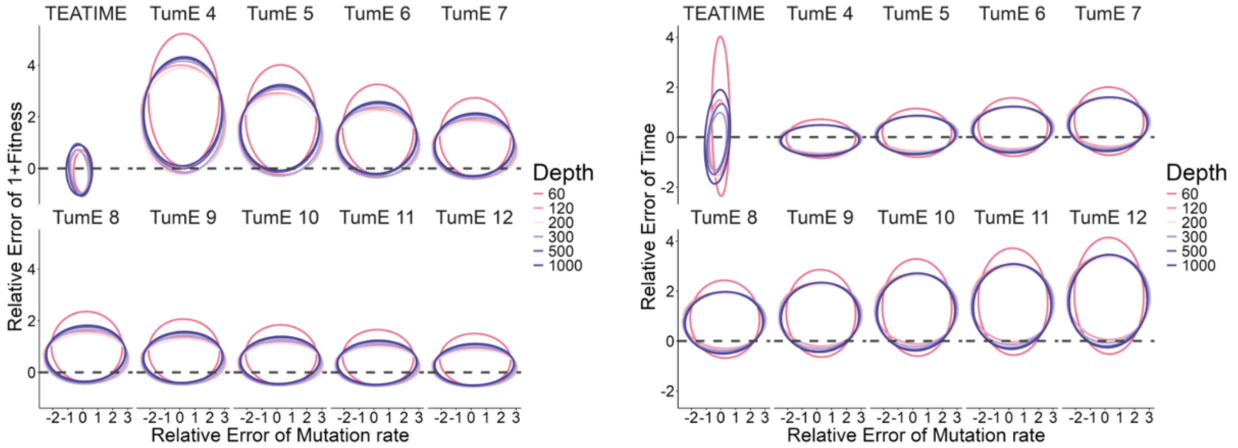


**Supplementary Figure 4.** Comparative analysis of the TEATIME method versus other models at population size N $\approx 10^6$ across different sequencing depths.

(A). Analyzable Ratio in different sequencing depths.

(B) Performance of MOBSTER using various N values, including $10^6$, compared to TEATIME.

(C) Performance of TumE (mean predictions) using various N values, including $10^6$, compared to TEATIME.

(D) 95% confidence ellipses showing the relative error distribution of parameter estimates

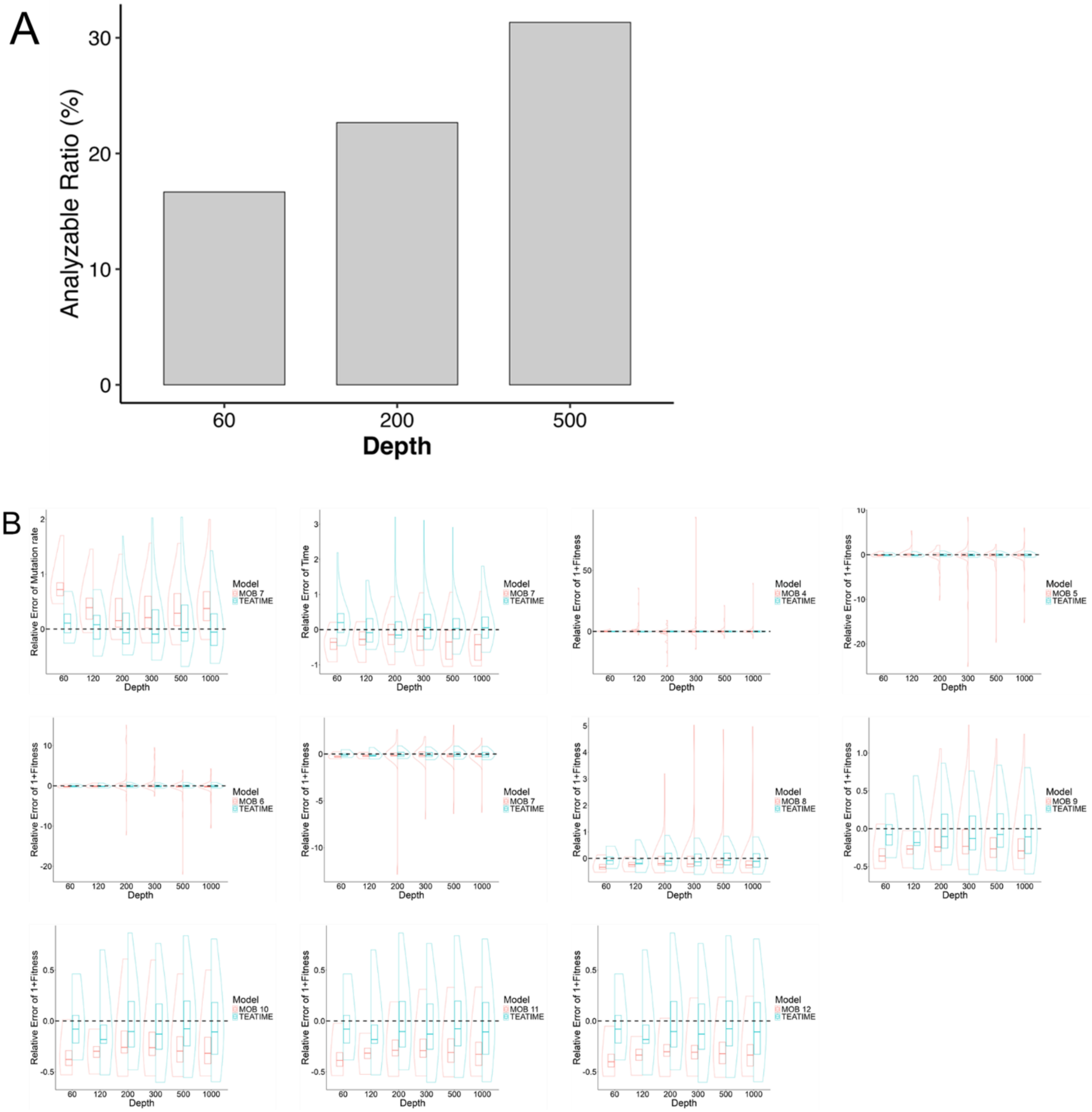
A
Analyzable Ratio (%)
0
10
20
30
60
200
500
Depth
B
Relative Error of Mutation rate
Relative Error of Time
Relative Error of 1+Fitness
Depth
Model
MOB 4
MOB 5
MOB 6
MOB 7
MOB 8
MOB 9
MOB 10
MOB 11
MOB 12
TEATIME
60
120
200
300
500
1000

C

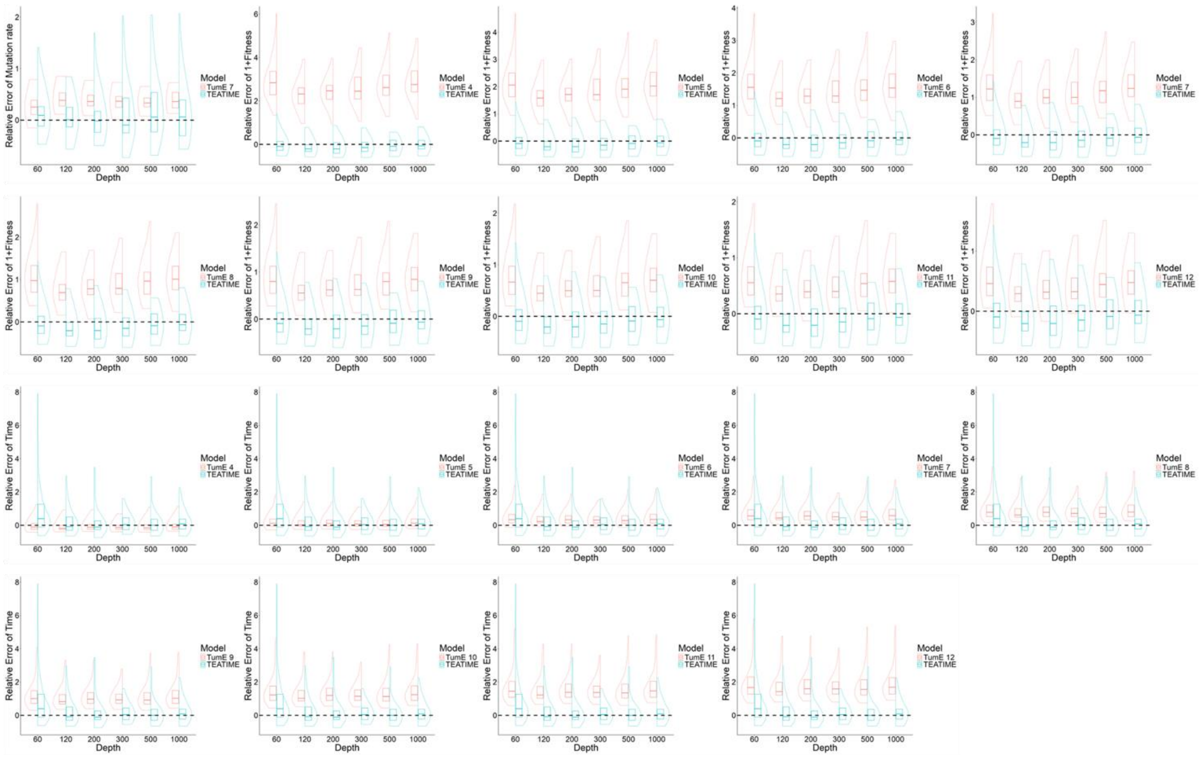

D

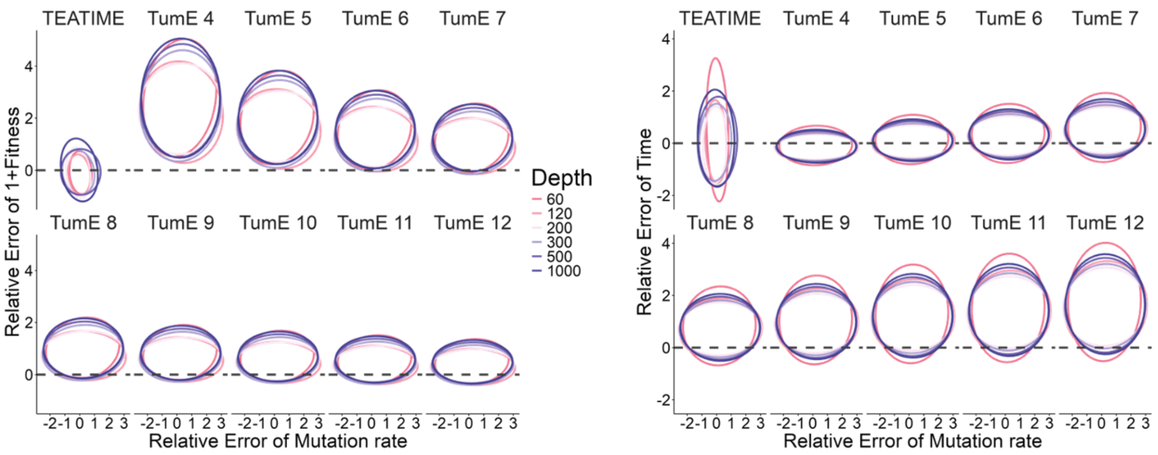


**Supplementary Figure 5.** Comparative analysis of the TEATIME method versus other models at population size N $\approx 10^7$ across different sequencing depths.

(A). Analyzable Ratio in different sequencing depths

(B) Performance of MOBSTER using various N values, including $10^7$, compared to TEATIME.

(C) Performance of TumE (mean predictions) using various N values, including $10^7$, compared to TEATIME.

(D) 95% confidence ellipses showing the relative error distribution of parameter estimates

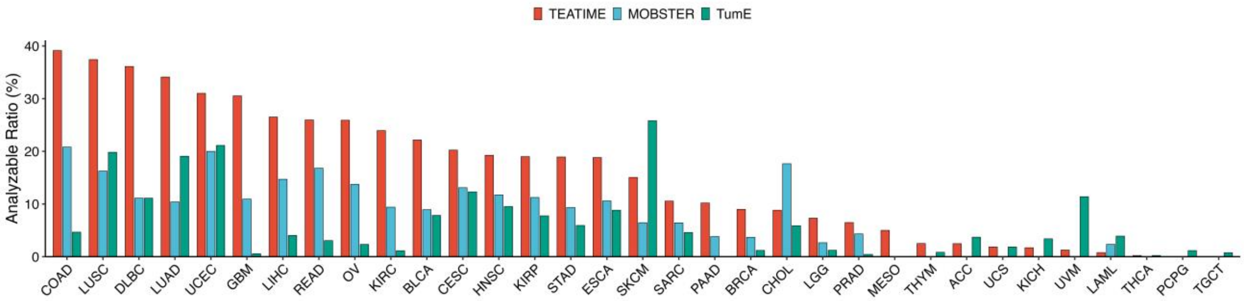


**Supplementary Figure 6.** Analyzable case ratio across tumor types in pan-cancer analysis. Tumor types are ordered by the proportion of predictable cases using TEATIME predictions.

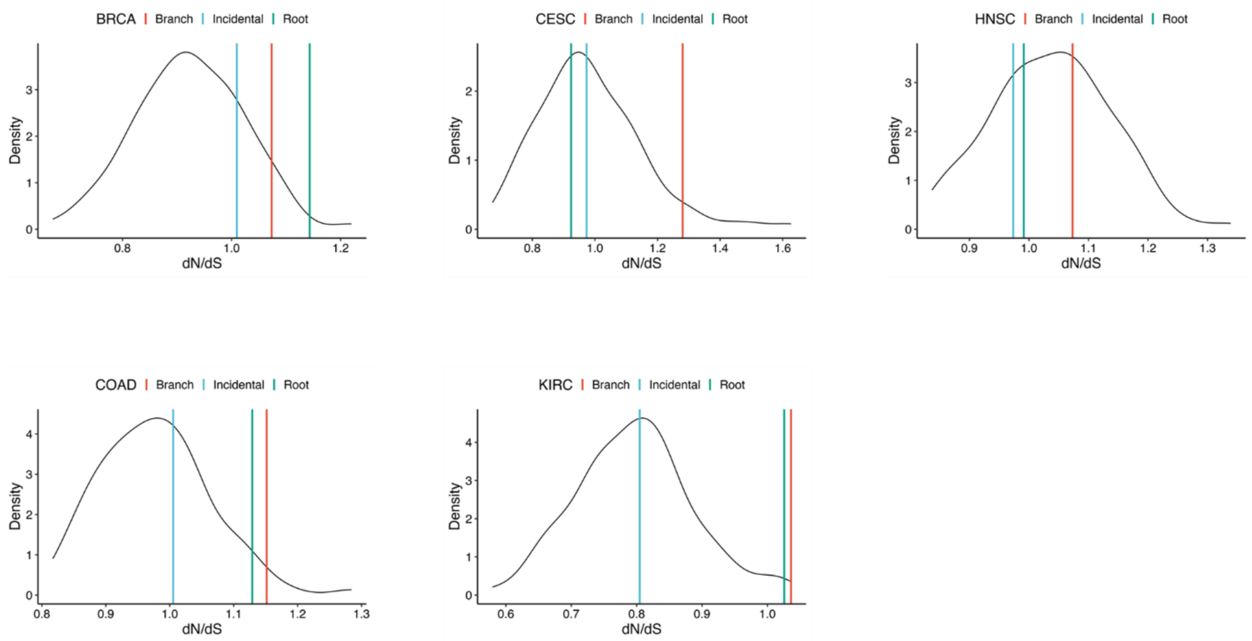


**Supplementary Figure 7.** dN/dS ratios in BRCA, COAD, CESC, HNSC, KIRC.

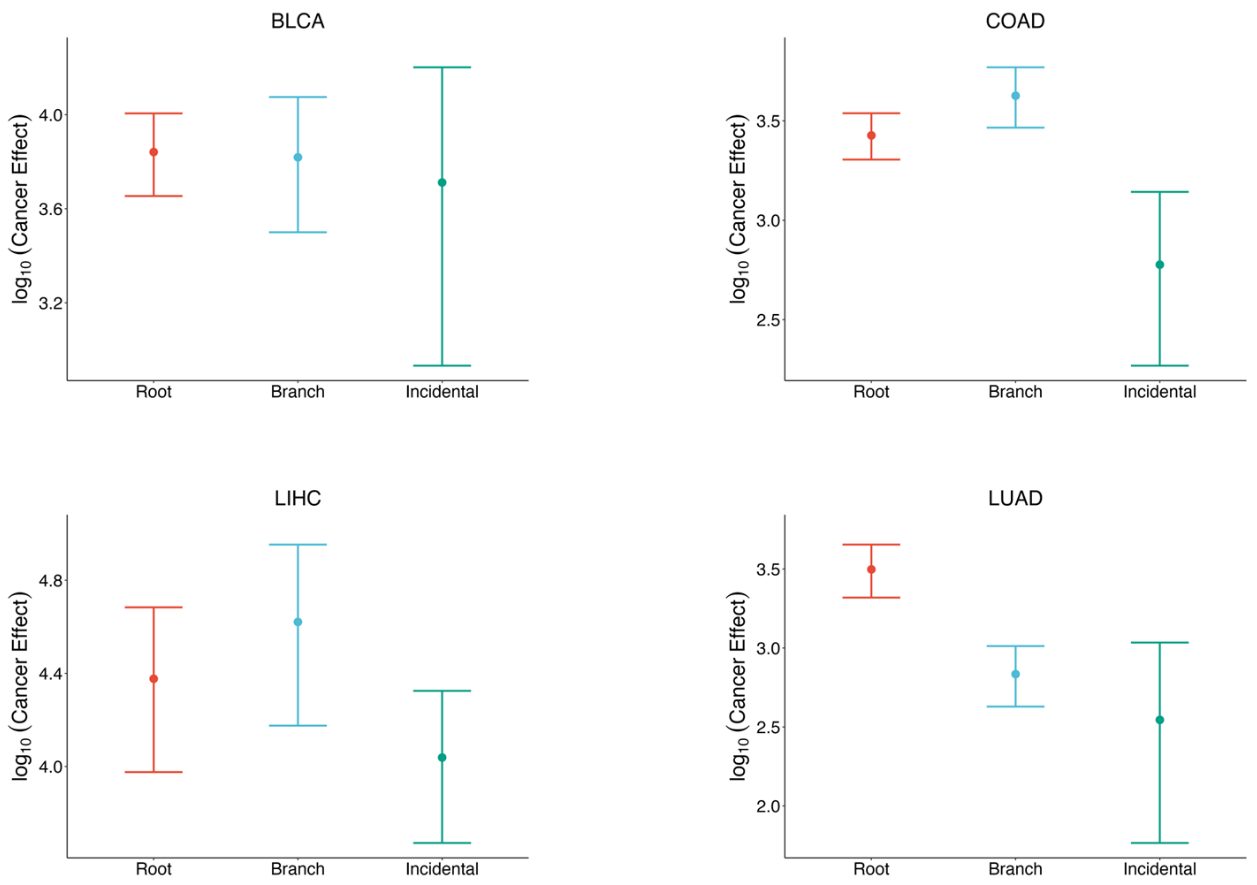


**Supplementary Figure 8.** Comparison of cancer effect size for Root, Branch and Incidental mutations across cancer types.

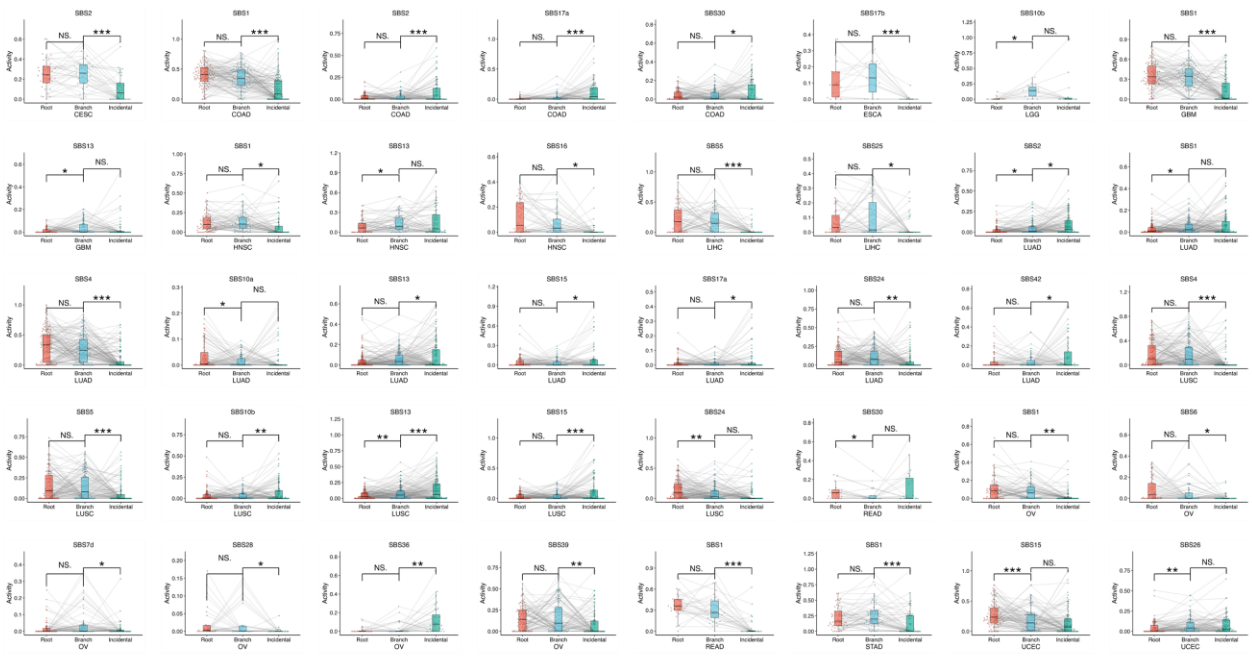

**Supplementary Figure 9.** Mutational signatures: Activity Changes moving from Root to Branch to Incidental mutations.

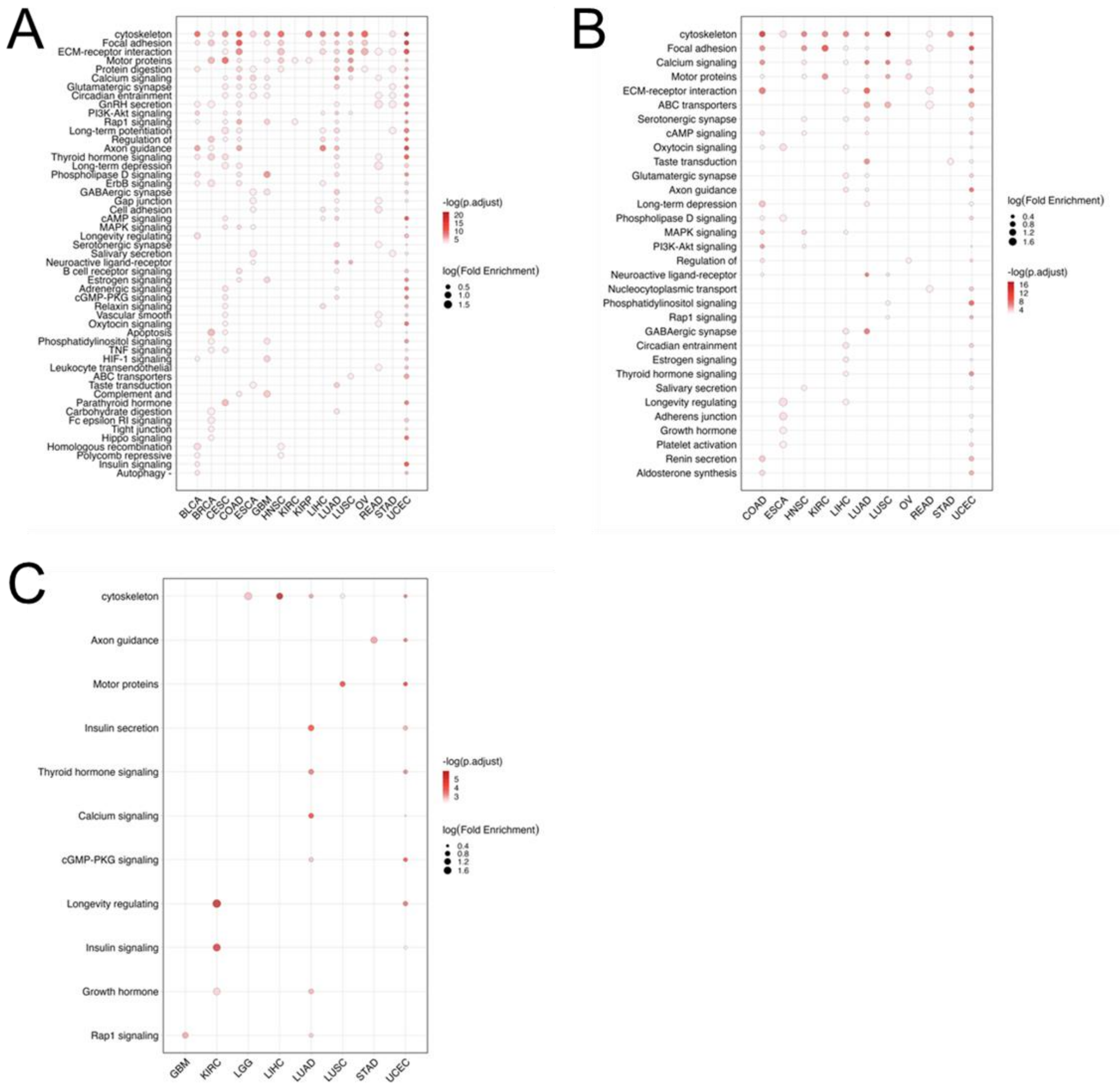


**Supplementary Figure 10.** Pan-cancer enrichment of pathways at Different Timepoint. (A) Enrichment on Root. (B) Enrichment on Branch. (C) Enrichment on Incidental. Only pathways significantly enriched in at least two cancer types are shown.

**Supplementary Table 1**. Primary tumor statistics.

| | | # of analyzable tumors | | |
|---|---|---|---|---|
| Cancer Type | # of primary tumors | MOBSTER | TEATIME | TUME |
| ACC | 81 | 0 | 2 | 3 |
| BLCA | 370 | 33 | 82 | 29 |
| BRCA | 923 | 34 | 83 | 11 |
| CESC | 252 | 33 | 51 | 31 |
| CHOL | 34 | 6 | 3 | 2 |
| COAD | 322 | 67 | 126 | 15 |
| DLBC | 36 | 4 | 13 | 4 |
| ESCA | 170 | 18 | 32 | 15 |
| GBM | 357 | 39 | 109 | 2 |
| HNSC | 452 | 53 | 87 | 43 |
| KICH | 59 | 0 | 1 | 2 |
| KIRC | 351 | 33 | 84 | 4 |
| KIRP | 258 | 29 | 49 | 20 |
| LAML | 128 | 3 | 1 | 5 |
| LGG | 492 | 13 | 36 | 6 |
| LIHC | 347 | 51 | 92 | 14 |
| LUAD | 519 | 54 | 177 | 99 |
| LUSC | 449 | 73 | 168 | 89 |
| MESO | 80 | 0 | 4 | 0 |
| OV | 386 | 53 | 100 | 9 |
| PAAD | 157 | 6 | 16 | 0 |
| PCPG | 173 | 0 | 0 | 2 |
| PRAD | 463 | 20 | 30 | 2 |
| READ | 131 | 22 | 34 | 4 |
| SARC | 218 | 14 | 23 | 10 |
| SKCM | 93 | 6 | 14 | 24 |
| STAD | 354 | 33 | 67 | 21 |
| TGCT | 131 | 0 | 0 | 1 |
| THCA | 451 | 0 | 1 | 1 |
| THYM | 120 | 0 | 3 | 1 |
| UCEC | 445 | 89 | 138 | 94 |
| UCS | 54 | 0 | 1 | 1 |
| UVM | 79 | 0 | 1 | 9 |

**Supplementary Table 2**. Mediation Test with immune infiltration

| Immune infiltration | value_type | Mediator | acme_estimate | acme_p | ade_estimate | ade_p | total_estimate | total_p | prop_estimate | prop_p |
|---|---|---|---|---|---|---|---|---|---|---|
| B.cell_TIMER | binary | Fitness | -0.033976657 | 0.016 | -0.16071075 | 0.018 | -0.1946874 | 0.006 | 0.1760557 | 0.022 |
| B.cell_TIMER | continuous | Fitness | -0.038598925 | 0.034 | -0.16067056 | 0.028 | -0.19926948 | 0.006 | 0.191354 | 0.04 |
| B.cell_XCELL | binary | Fitness | -0.043565368 | 0 | -0.27315125 | 0 | -0.31671662 | 0 | 0.13774345 | 0 |
| B.cell_XCELL | continuous | Fitness | -0.076189997 | 0 | -0.26122126 | 0 | -0.33741126 | 0 | 0.22459049 | 0 |
| B.cell.memory_CIBERSORT.ABS | binary | Fitness | -0.042957523 | 0.022 | -1.45005034 | 0 | -1.49300787 | 0 | 0.02950463 | 0.022 |
| B.cell.memory_CIBERSORT.ABS | continuous | Fitness | -0.198456926 | 0.032 | -1.44255654 | 0 | -1.64101347 | 0 | 0.11967468 | 0.032 |
| B.cell.memory_XCELL | binary | Fitness | -0.044740534 | 0 | -0.79817391 | 0 | -0.84291444 | 0 | 0.05338261 | 0 |
| B.cell.memory_XCELL | continuous | Fitness | -0.190042052 | 0 | -0.78569872 | 0 | -0.97574077 | 0 | 0.19335793 | 0 |
| B.cell.naive_XCELL | binary | Fitness | -0.045545378 | 0.002 | -1.54153938 | 0.012 | -1.58708476 | 0.012 | 0.02891556 | 0.014 |
| B.cell.naive_XCELL | continuous | Fitness | -0.429614341 | 0.01 | -1.56310343 | 0.01 | -1.99271777 | 0 | 0.21578302 | 0.01 |
| Cancer.associated.fibroblast_XCELL | binary | Fitness | -0.034525875 | 0.012 | -0.40837481 | 0 | -0.44290068 | 0 | 0.07993316 | 0.012 |
| Cancer.associated.fibroblast_XCELL | continuous | Fitness | -0.061493954 | 0.012 | -0.4176469 | 0 | -0.47914085 | 0 | 0.12750787 | 0.012 |
| Hematopoietic.stem.cell_XCELL | continuous | Fitness | -0.03258979 | 0.032 | -0.08460473 | 0.086 | -0.11719452 | 0.02 | 0.27326683 | 0.052 |
| immune.score_XCELL | binary | Fitness | -0.040932135 | 0 | -0.17167274 | 0 | -0.21260487 | 0 | 0.19128675 | 0 |
| immune.score_XCELL | continuous | Fitness | -0.068229166 | 0 | -0.15656165 | 0.002 | -0.22479082 | 0 | 0.30219096 | 0 |
| Macrophage_TIMER | continuous | Fitness | -0.057030504 | 0.006 | -0.10860026 | 0.146 | -0.16563076 | 0.034 | 0.33211836 | 0.04 |
| Macrophage_XCELL | binary | Fitness | -0.046650426 | 0.004 | -0.26789655 | 0.062 | -0.31454698 | 0.024 | 0.14811058 | 0.028 |
| Macrophage_XCELL | continuous | Fitness | -0.186409905 | 0 | -0.20610143 | 0.118 | -0.39251133 | 0.006 | 0.48290983 | 0.006 |
| Macrophage_XCELL | binary | Time | -0.010715692 | 0.03 | -0.37686664 | 0 | -0.38758233 | 0 | 0.02665683 | 0.03 |
| Macrophage.M1_CIBERSORT.ABS | binary | Fitness | -0.04690493 | 0 | -0.44772801 | 0.002 | -0.49463294 | 0 | 0.09527794 | 0 |
| Macrophage.M1_CIBERSORT.ABS | continuous | Fitness | -0.173655147 | 0 | -0.38686175 | 0.022 | -0.5605169 | 0 | 0.30672657 | 0 |
| Macrophage.M1_XCELL | binary | Fitness | -0.047461763 | 0.002 | -0.58554799 | 0.002 | -0.63300975 | 0.002 | 0.07603096 | 0.004 |
| Macrophage.M1_XCELL | continuous | Fitness | -0.283491016 | 0 | -0.49876685 | 0.018 | -0.78225786 | 0 | 0.35740113 | 0 |
| Macrophage.M1_XCELL | binary | Time | -0.012142612 | 0.026 | -0.73317137 | 0 | -0.74531398 | 0 | 0.01617303 | 0.026 |
| Macrophage.M2_CIBERSORT.ABS | continuous | Fitness | -0.067477249 | 0 | -0.06717743 | 0.298 | -0.13465468 | 0.042 | 0.48453298 | 0.042 |
| Macrophage.M2_XCELL | binary | Fitness | -0.046808203 | 0.018 | -0.50755653 | 0.038 | -0.55436474 | 0.028 | 0.08347552 | 0.046 |
| Macrophage.M2_XCELL | continuous | Fitness | -0.233219114 | 0.002 | -0.42296778 | 0.098 | -0.65618689 | 0.006 | 0.35398861 | 0.008 |
| Macrophage.M2_XCELL | binary | Time | -0.01376677 | 0.026 | -0.56277729 | 0.026 | -0.57654405 | 0.022 | 0.02353836 | 0.048 |
| microenvironment.score_XCELL | binary | Fitness | -0.041320321 | 0 | -0.18565454 | 0 | -0.22697486 | 0 | 0.18029796 | 0 |
| microenvironment.score_XCELL | continuous | Fitness | -0.065459483 | 0 | -0.17224883 | 0 | -0.23770832 | 0 | 0.27810751 | 0 |
| microenvironment.score_XCELL | binary | Time | -0.007558952 | 0.044 | -0.23430585 | 0 | -0.24186481 | 0 | 0.03065497 | 0.044 |
| Monocyte_XCELL | binary | Fitness | -0.045390058 | 0.002 | -0.44329524 | 0.004 | -0.4886853 | 0 | 0.09236373 | 0.002 |
| Monocyte_XCELL | continuous | Fitness | -0.19919794 | 0 | -0.38740315 | 0.016 | -0.58660109 | 0 | 0.33444572 | 0 |
| Monocyte_XCELL | binary | Time | -0.010965257 | 0.036 | -0.58325067 | 0 | -0.59421593 | 0 | 0.0183946 | 0.036 |
| Myeloid.dendritic.cell_TIMER | binary | Fitness | -0.024528784 | 0 | -0.08027054 | 0.004 | -0.10479933 | 0 | 0.23423425 | 0 |
| Myeloid.dendritic.cell_TIMER | continuous | Fitness | -0.036898163 | 0 | -0.06940229 | 0.006 | -0.10630045 | 0 | 0.3488855 | 0 |
| Myeloid.dendritic.cell_TIMER | binary | Time | -0.003826345 | 0.022 | -0.09651426 | 0.002 | -0.1003406 | 0.002 | 0.03574446 | 0.024 |
| Myeloid.dendritic.cell_XCELL | binary | Fitness | -0.04249724 | 0.038 | -0.51690563 | 0.008 | -0.55940287 | 0.002 | 0.07811728 | 0.04 |
| Myeloid.dendritic.cell_XCELL | continuous | Fitness | -0.167072359 | 0.012 | -0.46880502 | 0.024 | -0.63587738 | 0.004 | 0.26226418 | 0.016 |
| Myeloid.dendritic.cell.activated_XCEL | binary | Fitness | -0.039518015 | 0 | -0.16332781 | 0.002 | -0.20284582 | 0 | 0.19473056 | 0 |
| Myeloid.dendritic.cell.activated_XCEL | continuous | Fitness | -0.060291624 | 0 | -0.14748131 | 0 | -0.20777293 | 0 | 0.28993353 | 0 |
| Neutrophil_TIMER | binary | Fitness | -0.049379275 | 0 | -0.24454441 | 0.002 | -0.29392368 | 0 | 0.16983177 | 0 |
| Neutrophil_TIMER | continuous | Fitness | -0.114074208 | 0 | -0.20908076 | 0.01 | -0.32315497 | 0 | 0.35013614 | 0 |
| Neutrophil_TIMER | binary | Time | -0.008747558 | 0.03 | -0.30810004 | 0 | -0.3168476 | 0 | 0.02624185 | 0.03 |
| Plasmacytoid.dendritic.cell_XCELL | binary | Fitness | -0.047480882 | 0 | -0.30448238 | 0.042 | -0.35196326 | 0.018 | 0.13613797 | 0.018 |
| Plasmacytoid.dendritic.cell_XCELL | continuous | Fitness | -0.19338888 | 0 | -0.27288356 | 0.046 | -0.46627244 | 0 | 0.41699518 | 0 |
| stroma.score_XCELL | binary | Fitness | -0.040050483 | 0.014 | -0.25404321 | 0.008 | -0.2940937 | 0 | 0.13890811 | 0.014 |
| stroma.score_XCELL | continuous | Fitness | -0.080577364 | 0.01 | -0.24383436 | 0.014 | -0.32441173 | 0 | 0.24755821 | 0.01 |
| stroma.score_XCELL | binary | Time | -0.013678173 | 0.012 | -0.29198337 | 0 | -0.30566154 | 0 | 0.04467335 | 0.012 |
| T.cell.CD4._TIMER | continuous | Fitness | -0.038846557 | 0.024 | -0.11406221 | 0.066 | -0.15290876 | 0.014 | 0.25501937 | 0.038 |
| T.cell.CD4._TIMER | binary | Time | -0.00836564 | 0.04 | -0.13346134 | 0.042 | -0.14182697 | 0.028 | 0.05572147 | 0.064 |
| T.cell.CD4..naive_XCELL | binary | Fitness | -0.044653797 | 0.006 | -1.07095515 | 0.004 | -1.11560894 | 0.002 | 0.03995882 | 0.008 |
| T.cell.CD4..naive_XCELL | continuous | Fitness | -0.245364372 | 0.012 | -1.0660363 | 0 | -1.31140067 | 0 | 0.18686905 | 0.012 |
| T.cell.CD4..Th2_XCELL | binary | Fitness | -0.024597249 | 0.026 | -0.07284338 | 0.078 | -0.09744063 | 0.024 | 0.24270039 | 0.05 |
| T.cell.CD4..Th2_XCELL | continuous | Fitness | -0.050183825 | 0 | -0.05145613 | 0.2 | -0.10163996 | 0.016 | 0.48862705 | 0.016 |
| T.cell.CD8._CIBERSORT.ABS | binary | Fitness | -0.047861608 | 0 | -0.14831307 | 0.104 | -0.19617468 | 0.034 | 0.2399073 | 0.034 |
| T.cell.CD8._CIBERSORT.ABS | continuous | Fitness | -0.096059117 | 0 | -0.12489569 | 0.168 | -0.22095481 | 0.028 | 0.42349322 | 0.028 |
| T.cell.CD8._CIBERSORT.ABS | binary | Time | -0.009111001 | 0.036 | -0.2318176 | 0.006 | -0.2409286 | 0.004 | 0.03660359 | 0.04 |
| T.cell.CD8._TIMER | binary | Fitness | -0.040181893 | 0 | -0.06160205 | 0.102 | -0.10178395 | 0.012 | 0.40171411 | 0.012 |
| T.cell.CD8._TIMER | continuous | Fitness | -0.0563643 | 0 | -0.05537299 | 0.176 | -0.1117373 | 0.008 | 0.50843941 | 0.008 |
| T.cell.CD8._TIMER | binary | Time | -0.006383363 | 0.028 | -0.09092982 | 0.024 | -0.09731319 | 0.014 | 0.06266387 | 0.038 |
| T.cell.CD8._XCELL | binary | Time | -0.009334572 | 0.04 | -0.25476425 | 0.042 | -0.26409882 | 0.034 | 0.03328804 | 0.066 |
| T.cell.CD8..central.memory_XCELL | binary | Fitness | -0.045557383 | 0 | -0.18559481 | 0.038 | -0.2311522 | 0.008 | 0.19499893 | 0.008 |
| T.cell.CD8..central.memory_XCELL | continuous | Fitness | -0.096962242 | 0 | -0.17087923 | 0.044 | -0.26784147 | 0 | 0.36711751 | 0 |

**Supplementary Table 3**. Multivariable regression with immune infiltration

| method | Predictor | Immune infiltration | tumor | Standardized Estimate | pval_adj |
|---|---|---|---|---|---|
| TIMER | Diversity | B.cell_TIMER | SARC | -0.5434479 | 0.0500877 |
| TIMER | Fitness | B.cell_TIMER | HNSC | -0.308881 | 0.07253466 |
| TIMER | Fitness | B.cell_TIMER | LUSC | -0.2020764 | 0.04725699 |
| XCELL | Fitness | B.cell_XCELL | LUSC | -3.62E-01 | 0.00027253 |
| CIBERSORT | Diversity | B.cell.memory_CIBERSORT | LGG | -5.78E-01 | 0.01483937 |
| CIBERSORT | Time | B.cell.memory_CIBERSORT | LGG | 6.67E-01 | 0.00178503 |
| CIBERSORT | Diversity | B.cell.memory_CIBERSORT.ABS | LGG | -4.87E-01 | 0.07990545 |
| CIBERSORT | Time | B.cell.memory_CIBERSORT.ABS | LGG | 5.60E-01 | 0.02322822 |
| XCELL | Fitness | B.cell.memory_XCELL | LUSC | -3.30E-01 | 0.00090717 |
| CIBERSORT | Diversity | B.cell.naive_CIBERSORT.ABS | SARC | -5.77E-01 | 0.08517683 |
| XCELL | Time | B.cell.naive_XCELL | LUSC | 1.88E-01 | 0.09805491 |
| XCELL | Fitness | B.cell.plasma_XCELL | LUSC | -3.24E-01 | 0.00090717 |
| XCELL | Fitness | Cancer.associated.fibroblast_XCELL | LUSC | -2.17E-01 | 0.05030834 |
| XCELL | Diversity | Class.switched.memory.B.cell_XCELL | SARC | -5.67E-01 | 0.07940192 |
| XCELL | Fitness | Class.switched.memory.B.cell_XCELL | LUSC | -3.01E-01 | 0.00216981 |
| XCELL | Diversity | Common.lymphoid.progenitor_XCELL | LUSC | -1.94E-01 | 0.08644337 |
| XCELL | Fitness | Common.lymphoid.progenitor_XCELL | UCEC | -2.57E-01 | 0.08058452 |
| XCELL | Time | Endothelial.cell_XCELL | LUSC | 2.17E-01 | 0.05030834 |
| XCELL | Fitness | Eosinophil_XCELL | UCEC | 2.56E-01 | 0.08058452 |
| XCELL | Diversity | Granulocyte.monocyte.progenitor_XCELL | BRCA | -3.51E-01 | 0.0349398 |
| XCELL | Time | Granulocyte.monocyte.progenitor_XCELL | LUSC | 2.63E-01 | 0.00980389 |
| XCELL | Fitness | Hematopoietic.stem.cell_XCELL | LGG | -6.17E-01 | 0.03714285 |
| XCELL | Fitness | immune.score_XCELL | LUSC | -3.14E-01 | 0.00118415 |
| TIMER | Fitness | Macrophage_TIMER | BLCA | -0.2152666 | 0.0997433 |
| TIMER | Fitness | Macrophage_TIMER | LGG | -0.4600435 | 0.03511876 |
| TIMER | Time | Macrophage_TIMER | BLCA | 0.2643012 | 0.05268092 |
| CIBERSORT | Time | Macrophage.M1_CIBERSORT | UCEC | 2.73E-01 | 0.05422294 |
| CIBERSORT | Fitness | Macrophage.M1_CIBERSORT.ABS | LUSC | -3.05E-01 | 0.01169148 |
| XCELL | Fitness | Macrophage.M1_XCELL | LUSC | -2.36E-01 | 0.02912376 |
| XCELL | Fitness | Macrophage.M2_XCELL | LGG | -5.82E-01 | 0.02403316 |
| CIBERSORT | Diversity | Mast.cell.activated_CIBERSORT.ABS | SARC | -6.47E-01 | 0.08517683 |
| XCELL | Fitness | microenvironment.score_XCELL | LUSC | -3.27E-01 | 0.00090717 |
| XCELL | Fitness | Monocyte_XCELL | LUSC | -1.97E-01 | 0.07693762 |
| TIMER | Diversity | Myeloid.dendritic.cell_TIMER | BLCA | -0.268178 | 0.06304311 |
| TIMER | Fitness | Myeloid.dendritic.cell_TIMER | BLCA | -0.2299594 | 0.0986931 |
| TIMER | Fitness | Myeloid.dendritic.cell_TIMER | HNSC | -0.3627937 | 0.03311552 |
| TIMER | Fitness | Myeloid.dendritic.cell_TIMER | KIRC | -0.2612224 | 0.0915597 |
| TIMER | Fitness | Myeloid.dendritic.cell_TIMER | LUSC | -0.2321259 | 0.02515552 |
| TIMER | Time | Myeloid.dendritic.cell_TIMER | BLCA | 0.3460602 | 0.0213897 |
| TIMER | Time | Myeloid.dendritic.cell_TIMER | KIRC | 0.2502129 | 0.0915597 |
| XCELL | Diversity | Myeloid.dendritic.cell_XCELL | SARC | -6.43E-01 | 0.07940192 |
| XCELL | Fitness | Myeloid.dendritic.cell.activated_XCELL | LUSC | -2.87E-01 | 0.00369495 |
| CIBERSORT | Time | Myeloid.dendritic.cell.resting_CIBERSORT | LIHC | 4.44E-01 | 0.00274512 |
| CIBERSORT | Diversity | Myeloid.dendritic.cell.resting_CIBERSORT | KIRP | -4.82E-01 | 0.08799655 |
| CIBERSORT | Time | Myeloid.dendritic.cell.resting_CIBERSORT | LIHC | 5.01E-01 | 0.00032631 |
| CIBERSORT | Diversity | Neutrophil_CIBERSORT | KIRP | -4.61E-01 | 0.08799655 |
| CIBERSORT | Diversity | Neutrophil_CIBERSORT.ABS | KIRP | -0.4547739 | 0.08799655 |
| TIMER | Diversity | Neutrophil_TIMER | BLCA | -0.2568405 | 0.06304311 |
| TIMER | Fitness | Neutrophil_TIMER | BLCA | -0.2871608 | 0.05268092 |
| TIMER | Fitness | Neutrophil_TIMER | KIRC | -0.302086 | 0.0915597 |
| TIMER | Fitness | Neutrophil_TIMER | LUSC | -0.2916298 | 0.00350915 |
| TIMER | Time | Neutrophil_TIMER | BLCA | 0.2614325 | 0.05268092 |
| TIMER | Time | Neutrophil_TIMER | KIRC | 0.2469408 | 0.0915597 |
| CIBERSORT | Diversity | NK.cell.resting_CIBERSORT | LGG | -4.61E-01 | 0.09965729 |
| CIBERSORT | Diversity | NK.cell.resting_CIBERSORT.ABS | LGG | -4.87E-01 | 0.07990545 |
| XCELL | Fitness | Plasmacytoid.dendritic.cell_XCELL | LUSC | -2.39E-01 | 0.02673625 |
| XCELL | Fitness | stroma.score_XCELL | LUSC | -2.14E-01 | 0.05030834 |
| XCELL | Time | stroma.score_XCELL | LUSC | 2.16E-01 | 0.05030834 |
| TIMER | Diversity | T.cell.CD4._TIMER | BRCA | -0.289421 | 0.08199004 |
| TIMER | Diversity | T.cell.CD4._TIMER | SARC | -0.7550584 | 0.00683908 |
| TIMER | Fitness | T.cell.CD4._TIMER | BLCA | -0.2786026 | 0.05268092 |
| TIMER | Fitness | T.cell.CD4._TIMER | LUSC | -0.2252206 | 0.02515552 |
| XCELL | Diversity | T.cell.CD4...non.regulatory._XCELL | BRCA | -3.99E-01 | 0.0349398 |
| XCELL | Time | T.cell.CD4..central.memory_XCELL | BRCA | 3.35E-01 | 0.06019715 |
| XCELL | Diversity | T.cell.CD4..effector.memory_XCELL | SARC | -6.12E-01 | 0.07940192 |
| XCELL | Fitness | T.cell.CD4..effector.memory_XCELL | LUSC | -2.36E-01 | 0.02673625 |
| XCELL | Time | T.cell.CD4..effector.memory_XCELL | UCEC | 2.55E-01 | 0.08058452 |
| XCELL | Diversity | T.cell.CD4..memory_XCELL | SARC | -5.50E-01 | 0.07940192 |
| CIBERSORT | Diversity | T.cell.CD4..memory.resting_CIBERSORT.A | SARC | -0.727183 | 0.0264224 |
| XCELL | Diversity | T.cell.CD4..naive_XCELL | BRCA | -3.31E-01 | 0.06019715 |
| XCELL | Diversity | T.cell.CD4..naive_XCELL | SARC | -5.56E-01 | 0.07940192 |
| XCELL | Fitness | T.cell.CD4..naive_XCELL | LUSC | -2.05E-01 | 0.06181299 |
| XCELL | Time | T.cell.CD4..naive_XCELL | UCEC | 2.68E-01 | 0.08058452 |
| TIMER | Diversity | T.cell.CD8._TIMER | BRCA | -3.13E-01 | 0.08199004 |
| TIMER | Diversity | T.cell.CD8._TIMER | SARC | -5.20E-01 | 0.0500877 |
| TIMER | Time | T.cell.CD8._TIMER | BLCA | 0.3499961 | 0.0213897 |
| TIMER | Time | T.cell.CD8._TIMER | KIRC | 2.48E-01 | 0.0915597 |
| XCELL | Diversity | T.cell.CD8._XCELL | SARC | -5.57E-01 | 0.07940192 |
| XCELL | Fitness | T.cell.CD8._XCELL | LUSC | -1.99E-01 | 0.07693762 |
| XCELL | Diversity | T.cell.CD8..central.memory_XCELL | SARC | -5.89E-01 | 0.07940192 |
| XCELL | Fitness | T.cell.CD8..central.memory_XCELL | LUSC | -2.68E-01 | 0.00931648 |
| XCELL | Diversity | T.cell.CD8..naive_XCELL | BRCA | -3.69E-01 | 0.0349398 |
| CIBERSORT | Time | T.cell.follicular.helper_CIBERSORT | UCEC | 0.3796764 | 0.00073803 |
| CIBERSORT | Time | T.cell.follicular.helper_CIBERSORT.ABS | UCEC | 0.3048422 | 0.02433438 |
| XCELL | Diversity | T.cell.gamma.delta_XCELL | BRCA | -3.54E-01 | 0.0349398 |
| CIBERSORT | Diversity | T.cell.regulatory..Tregs._CIBERSORT.ABS | SARC | -0.618531 | 0.08517683 |
